# Highly recurrent multi-nucleotide mutations in SARS-CoV-2

**DOI:** 10.1101/2024.12.14.628346

**Authors:** Nicola De Maio, Kyle Smith, Yatish Turakhia, Nick Goldman

## Abstract

Multi-nucleotide mutations (MNMs) simultaneously replace multiple nucleotides. They are a significant contributor to evolution and disease, as well as to misdiagnosis, misannotation and other biases in genome data analysis.

MNMs are generally thought to be rare and random events. However, by processing millions of publicly shared genomes, we show that certain MNMs are highly recurrent in SARS-CoV-2: they repeatedly and consistently modify the same multiple nucleotides at the same genome position in the same way. The most frequent of these MNMs have independently occurred hundreds of times across all SARS-CoV-2 lineages.

The vast majority of these recurrent MNMs are linked to transcription regulatory sequences. We propose a mechanism that explains them through template switching as part of the natural transcription process of the virus.

This previously unknown mutational pattern increases our understanding of the evolution of SARS-CoV-2 and potentially many other nidoviruses. It also has important consequences for computational evolutionary biology: we show that for example recurrent MNMs cause approximately 14% of false positives during inference of recombination in SARS-CoV-2.

## 1 Introduction

Single-nucleotide substitutions are the most frequent type of genetic mutation[1], and are often the only ones considered in genome data analyses such as in population genetics[2] and phylogenetics[3]. However, other types of mutations also significantly contribute to genome evolution. Among these, multi-nucleotide mutations (MNMs), mutational events simultaneously replacing multiple nearby nucleotides, are some of the most debated[4–6].

Some early evidence for MNMs came from multi-nucleotide substitutions (MNSs: multiple nearby genetic differences between species)[4, 6–8]. Although the terms MNS and MNM are often used interchangeably, we define MNSs as nucleotide substitutions inferred on the same phylogenetic tree branch. A MNS could be the result of distinct sequential single-nucleotide mutation events, or of a single mutation event. On the other hand, we use the term MNM to refer to mutation events causing the instantaneous replacement of multiple nearby nucleotides. MNMs typically cause MNSs, but not all MNSs are attributable to MNMs, since successive near single-nucleotide mutations occurring on the same phylogenetic branch also cause MNSs.

It has been debated what proportion of MNSs (and specifically of MNSs comprising nearby nucleotide substitutions) is caused by MNMs, and what proportion might be caused by other factors such as selection[9, 10] and variation in substitution rates[5]. However, multiple lines of evidence such as trio sequencing and mutation accumulation lines have confirmed the existence of MNMs in different species, and have estimated their prevalence to be between 0.4% and 10% of total nucleotide mutations[11–17]. Causative mechanisms of MNMs include short-range template switches[18–20], and ultraviolet light DNA damage[12]. MNMs have been reported to be a significant source of disease in humans[21, 22] and other animals[23].

In computational biology, typically only single-nucleotide events are considered, causing problems like misannotation, and consequently mis-diagnosis[24, 25]. MNMs have also been shown to cause biases and false positives in positive selection inference[26–29]. For this reason, several authors have proposed the inclusion of MNMs in phylogenetic substitution models[6, 27–30].

The millions of SARS-COV-2 genomes published during the COVID-19 pandemic create the unprecedented opportunity to study the virus’s evolution with extraordinary power and detail[31–33]. Here, we use a highly curated dataset of >2 million SARS-CoV-2 genomes[34] to investigate MNMs. We take advantage of the low level of divergence and high density of sampling of this dataset to reconstruct the mutational history of the virus with near certainty, and of the large number of genomes to investigate site-specific mutational patterns with great detail and precision[31, 34]. Focusing on highly recurrent MNSs, we identify some as attributable to recurrent sequence artefacts but conclude that the majority are caused by recurrent MNMs. Some of these have occurred hundreds of times during SARS-CoV-2 evolution as identical but distinct mutation events. We find strong evidence that recurrent MNMs are caused by interrupted template switching, and we show that highly recurrent MNMs cause artefacts in the inference of recombination.

## 2 Results

We investigated a mutation-annotated tree (a phylogenetic tree enriched with the information of which mutation was estimated to have occurred on which branch) inferred from a public dataset of *>* 2 millions SARS-CoV-2 genomes[34, 35] (see Methods section). We consider an MNS to be any group of nucleotide substitutions occurring on the same branch, and do not impose any cutoff based on the location along the genome of the substitutions involved (we also keep track of groups of substitutions far from each other along the genome). Defining recurrence as the number of times a substitution occurs along the phylogenetic tree (not the number of genomes containing the substitutions), we list the most highly recurrent MNSs in Table 1. The genome positions of all substitutions considered here are represented with respect to the reference genome MN908947.3. The most recurrent MNSs found were A28877T-G28878C, which was inferred to have occurred 480 times, and was observed in 14,714 genomes; and C21304A-G21305A, inferred to have arisen 736 times: 452 times on its own (and observed in 5,330 genomes), and 284 separate times as part of the larger MNS C21302T-C21304A-G21305A (observed in 4,197 genomes).

**Table 1:** Highly recurrent MNSs. Those unlikely to be caused by recurrent MNMs are highlighted in red. Number of occurrences: number of times the MNS is inferred to have arisen. Occurrences of larger MNSs are not considered when counting occurrences of smaller MNSs. For example, occurrences of C21302T-C21304A-G21305A do not add to the count shown for C21304A-G21305A, although the latter is contained in the former. Individual occurrences: numbers of inferred substitutions for each single-nucleotide substitution composing the MNS, without counting occurrences of the MNS itself; for example, the first row states that 219 C21302T substitutions were inferred. Proportion of singletons: proportions of MNS events with exactly one descendant. Number of observed genomes: number of genomes in which the mutated version of the MNS is observed. Observed genomes per occurrence: value in column 5 divided by the one in column 2, that is, the mean number of descendants of a MNS event. Proposed mechanism: proposed cause of the recurrent MNS (in red when cause is not an MNM). Protein affected: protein affected by the MNM and type of change (omitted for non-MNM MNSs). ORF9c is in parentheses since it is not clear if it is functionally expressed[36].

| MNS | Number of occurrences | Individual occurrences | Proportion of singletons | Number of observed genomes | Observed genomes per occurrence | Proposed mechanism | Protein affected |
| --- | --- | --- | --- | --- | --- | --- | --- |
| C21302T<br>-C21304A<br>-G21305A | 284 | 219<br>369<br>290 | 0.59 | 4197 | 14.8 | plausible interrupted template switching | ORF1b: P2612L<br>ORF1b: R2613N |
| C21304A<br>-G21305A | 452 | 369<br>290 | 0.63 | 5330 | 11.8 | plausible interrupted template switching | ORF1b: R2613N |
| A28877T<br>-G28878C | 480 | 50<br>18 | 0.62 | 14714 | 30.7 | interrupted template switching | N: synonymous<br>(ORF9c: V49L) |
| G27382C<br>-A27383T<br>-T27384C | 253 | 135<br>75<br>1724 | 0.66 | 141947 | 561 | interrupted template switching | ORF6: D61L |
| T26491C<br>-A26492T<br>-T26497C | 160 | 21<br>36<br>137 | 0.72 | 351 | 2.2 | plausible interrupted template switching | (intergenic) |
| G27758A<br>-T27760A | 126 | 43<br>60 | 0.64 | 312 | 2.5 | plausible interrupted template switching | ORF7a: synonymous<br>ORF7b: M1I, I2N |
| T27875C<br>-C27881T<br>-G27882C<br>-C27883T | 49 | 33<br>470<br>25<br>198 | 0.55 | 1204 | 24.6 | interrupted template switching | ORF7b: frameshift |
| C27881T<br>-G27882C<br>-C27883T | 61 | 470<br>25<br>198 | 0.64 | 442 | 7.2 | interrupted template switching | ORF7b: frameshift |
| C25162A<br>-C25163A | 116 | 21<br>106 | 0.60 | 624 | 5.4 | plausible interrupted template switching | S: Q1201K |
| T21294A<br>-G21295A<br>-G21296A | 89 | 18<br>49<br>58 | 0.58 | 444 | 5.0 | plausible interrupted template switching | ORF1b: G2610N |
| A27038T<br>-T27039A<br>-C27040A | 87 | 7<br>17<br>10 | 0.83 | 195 | 2.2 | interrupted template switching | M: S173K |
| A507T<br>-T508C<br>-G509A | 89 | 7<br>69<br>73 | 0.82 | 166 | 1.9 | sequence error due to incomplete read trimming |  |
| T28881A<br>-G28882A<br>-G28883C | 59 | 19<br>42<br>19 | 0.58 | 848359 | 14379 | sequence errors due to reference bias |  |
| A21550C<br>-A21551T | 68 | 14<br>43 | 0.72 | 202 | 3.0 | interrupted template switching | ORF1b: N2694 -<br>ORF1b: N2695L |
| T21994C<br>-T21995C | 64 | 55<br>139 | 0.67 | 103 | 1.6 | sequence errors due to deletion |  |
| C13423A<br>-C13424A | 62 | 16<br>165 | 0.69 | 157 | 2.5 | plausible interrupted template switching | ORF1a: R4387S |
| A4576T<br>-T4579A | 60 | 22<br>533 | 0.62 | 2330 | 38.8 | plausible interrupted template switching | ORF1a: synonymous |
| A20284T<br>-T20285C | 57 | 1<br>7 | 0.61 | 138 | 2.4 | unknown MNM | ORF1b: I2273S |
| G11083T<br>-C21575T | 53 | 11206<br>6913 | 0.81 | 826 | 15.6 | successive single-nuc. substitutions |  |

We consider four possible causes of highly recurrent MNSs:

- Highly recurrent MNMs (Fig. 1A). While previous studies have confirmed the occurrence of MNMs in different organisms[11–17], to our knowledge, highly recurrent MNMs have not been previously reported; as such, we only consider recurrent MNMs as a plausible cause of recurrent MNSs if we can reasonably exclude all other causes below.
- Combinations of successive individual single-nucleotide mutations. Some sites of the SARS-CoV-2 genome are affected by extremely highly recurrent single-nucleotide mutations[31, 34]. If two highly recurrent single-nucleotide mutation events independently occur on the same branch of the phylogenetic tree, they cause a MNS (Fig. 1B). If this occurs often enough, the corresponding MNS becomes highly recurrent. This scenario is discussed in section 2.1.

**Figure 1:**
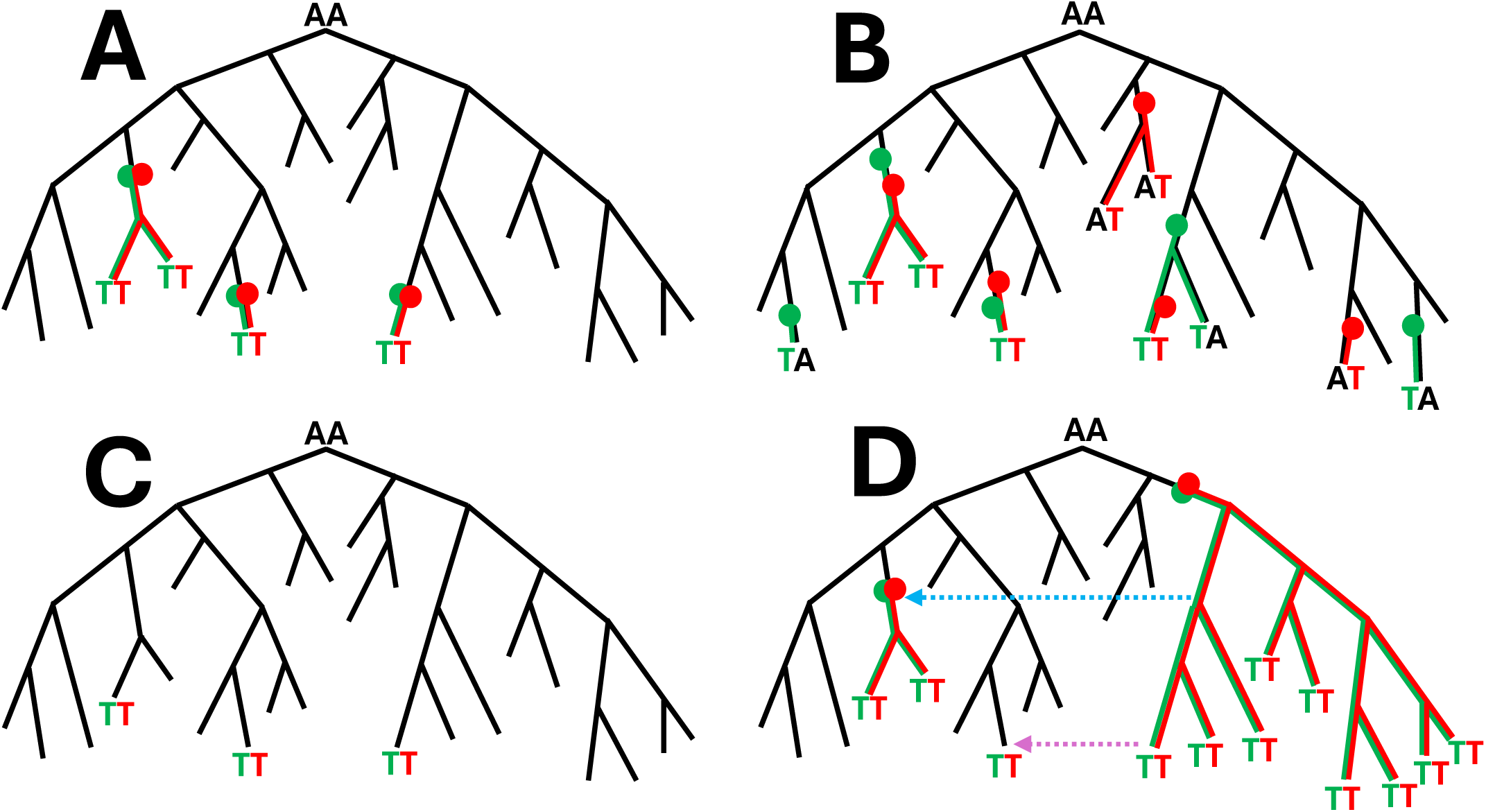
Graphical examples of causes of recurrent MNSs. We show a very small, illustrative phylogenetic tree in black. In all examples we show the substitution history of two genome positions for both of which the root nucleotide is A. Substitutions for the two positions are highlighted respectively in green and red. Substitution events along the tree are highlighted with colored circles, their descendants are highlighted with colored branches, and sequenced nucleotides different from the root “AA” are shown at the tips of the tree. **A** Recurrent MNMs: example recurrent AA to TT MNM. **B** Combinations of individual recurrent single-nucleotide mutations: A to T mutations at the two positions occur frequently and independently of each other; by chance, both mutations can occur on the same branch, giving rise to an AA to TT MNS that is not caused by an AA to TT MNM. **C** Consensus sequence and alignment artefact: errors at both genome positions can be correlated, for example caused by the same alignment error, resulting in AA to TT MNSs, at terminal tree branches only, that are not caused by AA to TT MNMs. **D** Recombination, contamination and mixed infections: when a MNS has many descendants (right side of the tree), recombinations (blue dashed arrow) that transfer the MNS into a different genetic background are more likely. Contamination or mixed infections (purple dashed arrow) can cause a similar artefactual pattern at terminal tree branches.

- Consensus sequence and alignment artefacts. Imperfections in sequencing and bioinformatics protocols can lead to recurrent errors in SARS-CoV-2 genome sequences and alignment[34, 37, 38]. Some of these problems can simultaneously affect more than one nucleotide, and lead to recurrent MNSs in inferred mutation-annotated trees[34, 37, 38] (Fig. 1C). In section 2.2 we discuss how we identify such artefacts by jointly analyzing sequencing read data and phylogenetic trees.
- Recombination and contamination. Recombination causes the exchange of part of the genome between different lineages[38, 39]. For recombination to occur in SARS-CoV-2, two different viral lineages need to infect the same host at the same time, and this is more likely when the two lineages are both circulating at high prevalence in the host population. If one, but not the other lineage contains a pre-existing MNS (either due to a MNM or a combination of successive mutations), then recombination can cause a new occurrence of the same MNS (Fig. 1D). If this occurs often enough, it might cause a highly recurrent MNS. Contamination (multiple lineages accidentally mixed in the same sample)[40] and mixed infections[41] can cause a similar observed pattern: if the two lineages present in the sample have varying sequencing read depth along the genome, and especially if an amplicon dropout affects only some of the two lineages, it can result in a mixed consensus sequence, that effectively looks like a recombinant (Fig. 1D). Recombination and contamination are discussed in section 2.3.

In Table 1, recurrent MNSs that we find unlikely to be caused by genuine recurrent MNMs are highlighted in red. Each is discussed below.

### 2.1 Combinations of highly recurrent single-nucleotide mutations

Substitution rate variation along the genome can increase the number of expected MNSs[5]. In SARS-CoV-2, extremely highly recurrent single-nucleotide mutations[31, 34] are indeed expected to cause recurrent MNSs when pairs of these mutations occur by chance on the same phylogenetic tree branch (Fig. 1B).

In our dataset, the most highly recurrent single-nucleotide substitutions are G11083T (11,206 substitutions) and C21575T (6,913 substitutions); their combination, G11083T-C21575T, with 53 occurrences, is the most recurrent MNS composed of non-neighbouring substitutions (Table 1). It is reasonable to conclude that occurrences of G11083T-C21575T are likely combinations of independently occurred single-nucleotide mutation events. For this reason, from here onward we only focus on MNSs more frequent than G11083T-C21575T and we exclude this MNS from consideration as a possible recurrent MNM.

All these remaining recurrent MNSs are composed of much rarer single-nucleotide substitutions than G11083T or C21575T (column “Individual substitutions” in Table 1). In fact, these MNSs are often more recurrent than the individual single-nucleotide substitutions they are composed of, and often involve more than two nucleotide substitutions. These MNSs are also more recurrent than the vast majority of single-nucleotide substitutions in the SARS-CoV-2 genome (Fig. 2A), with C21304A-G21305A and A28877T-G28878C being among the most highly recurrent (either single-nucleotide or multi-nucleotide) substitutions overall. It is therefore extremely unlikely that any substantial proportion of the occurrences of these remaining MNSs is the result of independently occurring single-nucleotide mutation events.

We did not investigate MNSs less recurrent than G11083T-C21575T, of which the most recurrent are C22716A-T22717C (45 occurrences), 27672A-C27673A (37 occurrences), and G910A-T911A-C912A (36 occurrences).

**Figure 2:**
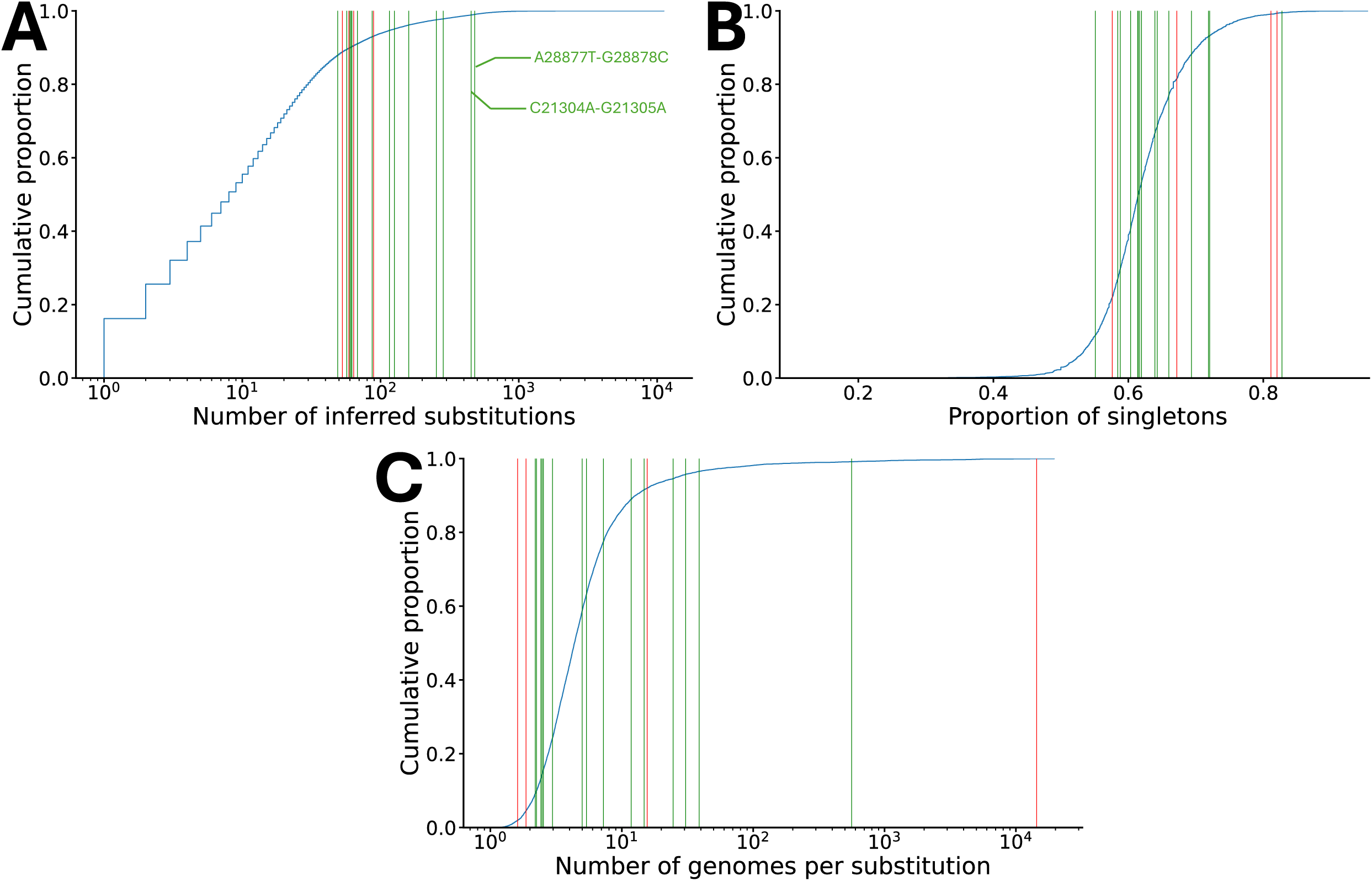
Comparison between recurrent MNSs and single-nucleotide substitutions in SARS-CoV-2. We show in blue the cumulative distributions for single-nucleotide substitutions. For any genome position, only the three substitutions away from the reference nucleotide are considered, and of these only those that are inferred to have occurred at least once (57,228 substitutions out of all possible 89,709). Vertical bars correspond to MNSs in Table 1: red bars correspond to those unlikely to be caused by MNMs (red entries in Table 1), green bars to all other MNSs. **A** Numbers of substitutions, as in Table 1. MNSs in Table 1 are more recurrent than most single-nucleotide substitutions, since the left-most green bar (corresponding to MNS T27875C-C27881T-G27882C-C27883T, which has 49 substitutions, see *x*-axis) intersects the blue cumulative distribution at a value of around 0.88, meaning that the least recurrent MNS in Table 1 is more recurrent than 88% of single-nucleotide substitutions observed in our dataset. **B** Proportions of substitutions that are singletons (i.e. occurring on terminal branches of the phylogenetic tree and therefore inherited by only one sampled genome), like column “Proportion of singletons” in Table 1. Substitutions due to recurrent sequence errors are typically enriched in singletons (they are represented at the right-hand side of the plot), as is the case for the right-most red vertical bar in the plot corresponding to MNS A507T-T508C-G509A. **C** Number of observed genomes containing the considered substitution, divided by the number of occurrences of that substitution along the phylogenetic tree; that is, the average number of sampled descendants of a substitution event (like column “Observed genomes per occurrence” in Table 1). Substitutions due to recurrent sequence errors are expected to have few genomes per substitution, while those with many descendant genomes per substitution might be more affected by reference biases and recombination.

### 2.2 Sequence and alignment artefacts

Artefacts in consensus genome calling and in alignments can cause MNSs[34, 37, 38]. Following [34], we use a combination of three approaches to identify instances of these issues in our SARS-CoV-2 data set:

- We investigate the proportions of singletons (substitutions occurring on terminal branches of the phylogenetic tree) for different types of substitution (section 2.2.1). While a true mutation can be inherited by many descendants, a consensus calling error only affects one sample at the time (see Fig. 1C), usually resulting in an enrichment of singleton substitutions at positions with recurrent sequence errors[34]. An enrichment in singletons is however not sufficient on its own to assess that a recurrent MNS is due to a recurrent error, since deleterious mutations are also expected to show an enrichment in singletons.
- We analyze sequencing read data (section 2.2.2). Recurrent sequence errors are often associated with low sequencing depth, high heterozygosity in the raw sequencing reads, and indels[34]. For each recurrent MNS considered, we systematically analyze sequencing read data of “cherries” (sibling pairs of samples in the phylogenetic tree) in which only one of the two siblings contains the MNS. In these situations, the sibling not containing the MNS shows the genomic background in which the MNS has occurred. For each such cherry, we investigate sequencing reads that Viridian[35] maps near the considered MNS. Sequencing read data from cherries can highlight for example if an MNS can be explained by read alignment errors near a pre-existing indel, by reference biases, or by incomplete primer trimming.
- We investigate phylogenetic clustering (section 2.2.3). A given recurrent error can preferentially appear in certain parts of the phylogenetic tree if, for example, it is caused by a particular sequencing approach or genomic background[34, 37, 38]. We therefore consider phylogenetic clustering as a red flag for sequence errors; we do not consider it as a sufficient condition to rule an MNS as artefactual, however, since phylogenetic clustering can also be caused by context-dependent mutational events.

#### 2.2.2 Proportion of singletons

Only three of the recurrent MNSs considered present unusually high proportions of singletons (Fig. 2B and Table 1): G11083T-C21575T (which we already discarded above), A507T-T508C-G509A, and A27038T-T27039A-C27040A. MNS A507T-T508C-G509A is indeed typically found in regions of sudden drop in sequencing depth (Supplementary Table S1) and appears to be the result of consensus sequence calling errors due to incomplete read trimming. As such, we also consider A507T-T508C-G509A to be problematic and unlikely the result of recurrent MNMs.

The MNS A27038T-T27039A-C27040A is also substantially enriched in singletons, but we found no other evidence to suggest that it is caused by artefacts (see Sections 2.2.2 and 2.2.3). For this reason, it is possible that A27038T-T27039A-C27040A (which causes an amino acid substitution in gene M, see Table 1 and Fig. 7) is caused by a genuine but deleterious recurrent MNM, reducing the ability of the virus to spread and infect new hosts.

#### 2.2.3 Read data for cherries

Recurrent heterozygosity in read data is often the hallmark of recurrent sequence errors[34, 37, 38], but apart from A507T-T508C-G509A and G11083T-C21575T which we already identified as problematic, no other MNS is highly enriched in heterozygosity in sequencing read data (column “Heterozygosity in cherries” in Table S2).

Recurrent consensus sequence errors can also emerge in regions of low read depth (e.g. due to systematic amplicon dropout). T21994C-T21995C is the only recurrent MNS appearing to be enriched in regions of reduced sequencing depth (*<* 100 in 26/28 cherries considered, see column “Low coverage in cherries” in Table S2). Indeed, T21994C-T21995C appears to be an artefact almost entirely specific to Alpha genomes (Supplementary Fig. S1A) caused by a pre-existing common deletion at positions 21991-21993, local sequence similarity (Supplementary Fig. S1B), and drops in sequencing depth (Supplementary Fig. S1C), causing read alignment and consensus calling errors.

T28881A-G28882A-G28883C often appears within a genomic background containing a deletion at these positions. While we avoid most reference biases in consensus sequence calls by using Viridian[35] (in particular, we prevent calling reference nucleotides at positions with very low coverage), we re-introduce some of this bias by masking short deletions (in this case, replacing gap characters with reference genome nucleotides) to improve phylogenetic inference[34]. Most inferred substitution events of T28881A-G28882A-G28883C, which is the most prevalent MNS among sampled genomes (Table 1), appear to be caused by this reference bias. T28881A-G28882A-G28883C seems to be the only recurrent MNS affected by this problem, since, due to its abundance, the nucleotides AAC at positions 28881-28883 are the genome-alignment-wide consensus nucleotides used for masking short deletions. In the following we consider this MNS as problematic and unlikely to be caused by recurrent MNMs.

None of the remaining MNSs systematically occurs near a pre-existing background indel. Instead, MNSs T26491C-A26492T-T26497C, T27875C-C27881T-G27882C-C27883T, C27881T-G27882C-C27883T, and A21550C-A21551T appear to consistently introduce certain indels (column “Deletions in cherries” in Table S2), specifically 1–3 nucleotide deletions, not present in the genomic background on which they occur (Supplementary Fig. S2). Since we find no other evidence to suggest these MNSs are artefactual, we suggest that these MNSs and deletions are the result of complex MNM events that not only substitute 2–4 nucleotides, but also consistently shorten the genome by 1–3 nucleotides.

#### 2.2.4 Phylogenetic clustering

Phylogenetic clustering of substitution events can be a red flag for recurrent sequence errors (for example highlighting errors caused by certain sequencing protocols[34]), but it can also emerge naturally as mutation rates might strongly depend on the background nucleotide context[31, 34]. As discussed in section 2.2.2, phylogenetic clustering is observed for the problematic MNS T21994C-T21995C (Supplementary Fig. S1A), linked to the presence of a background deletion causing read alignment errors.

We only observe obvious phylogenetic clustering for two other MNSs, T27875C-C27881T-G27882C-C27883T (Fig. 3) and A28877T-G28878C (Fig. 4; the phylogenetic distribution of the other MNSs is shown in Supplementary Fig. S3). Occurrences of T27875C-C27881T-G27882C-C27883T seem to particularly appear in the background of the nearby substitution C27874T (Fig. 3). Similarly, A28877T-G28878C appears to preferentially occur in the mutated background containing the nearby substitutions T28881A-G28882A-G28883C (Fig. 4). Unlike for T21994C-T21995C, we find no other evidence suggesting that T27875C-C27881T-G27882C-C27883T and A28877T-G28878C are not caused by genuine MNMs. This suggests that MNMs causing these MNSs might be preferentially happening in a specific nucleotide context; this is further explored in section 2.4.

**Figure 3:**
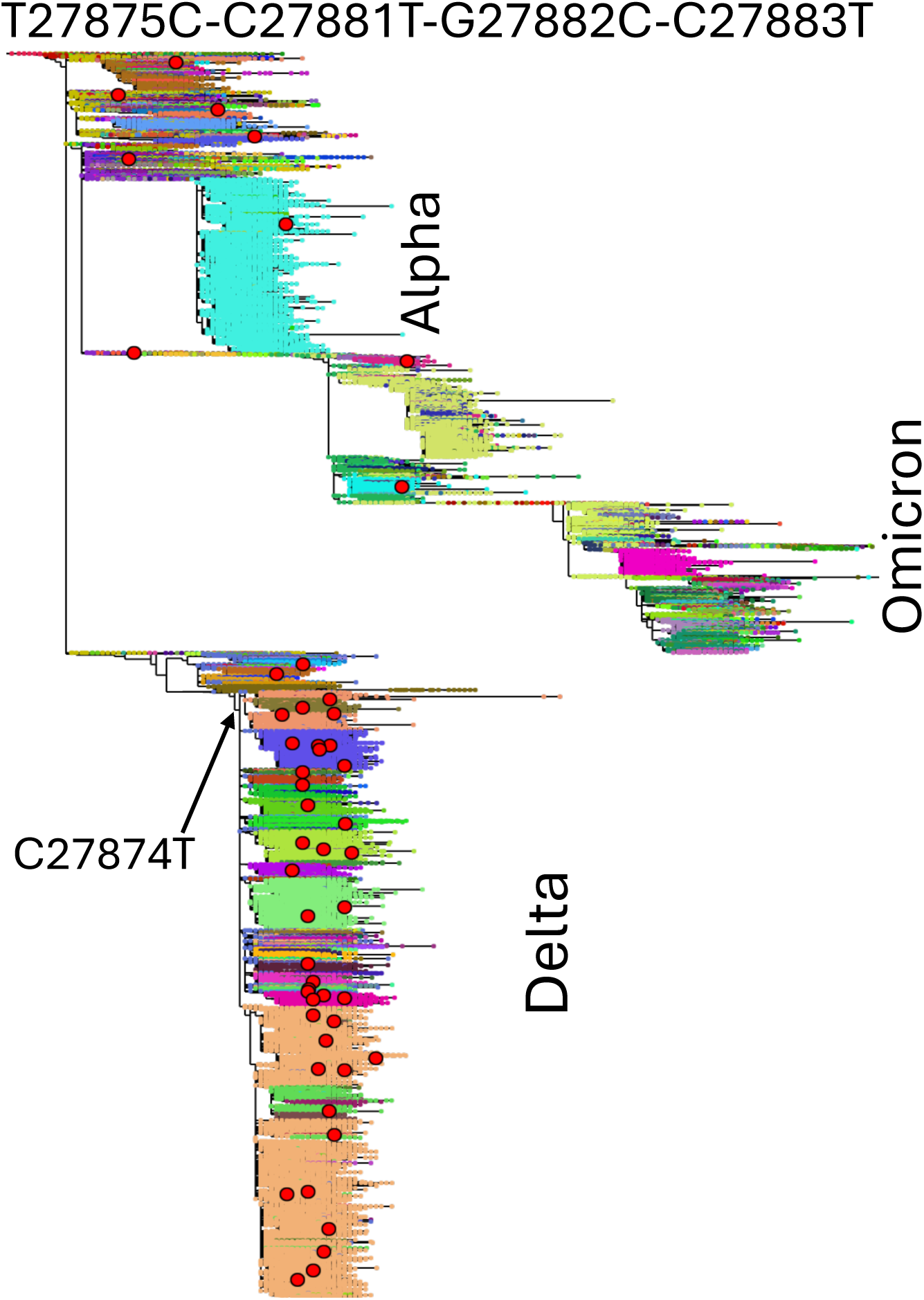
Phylogenetic distribution of MNS T27875C-C27881T-G27882C-C27883T. We show our background SARS-CoV-2 phylogenetic tree relating *>* 2 million genomes[34] where sampled genomes, represented by small dots, are colored according to their Pango[42] lineage (assigned by Pangolin[43] V4.3 with pangolin-data V1.21). Occurrences of T27875C-C27881T-G27882C-C27883T are highlighted with red circles. These appear enriched in the Delta lineage, where substitution C27874T (arrowed) is ancestral to most genomes. The tree and substitutions are visualized with Taxonium[44].

**Figure 4:**
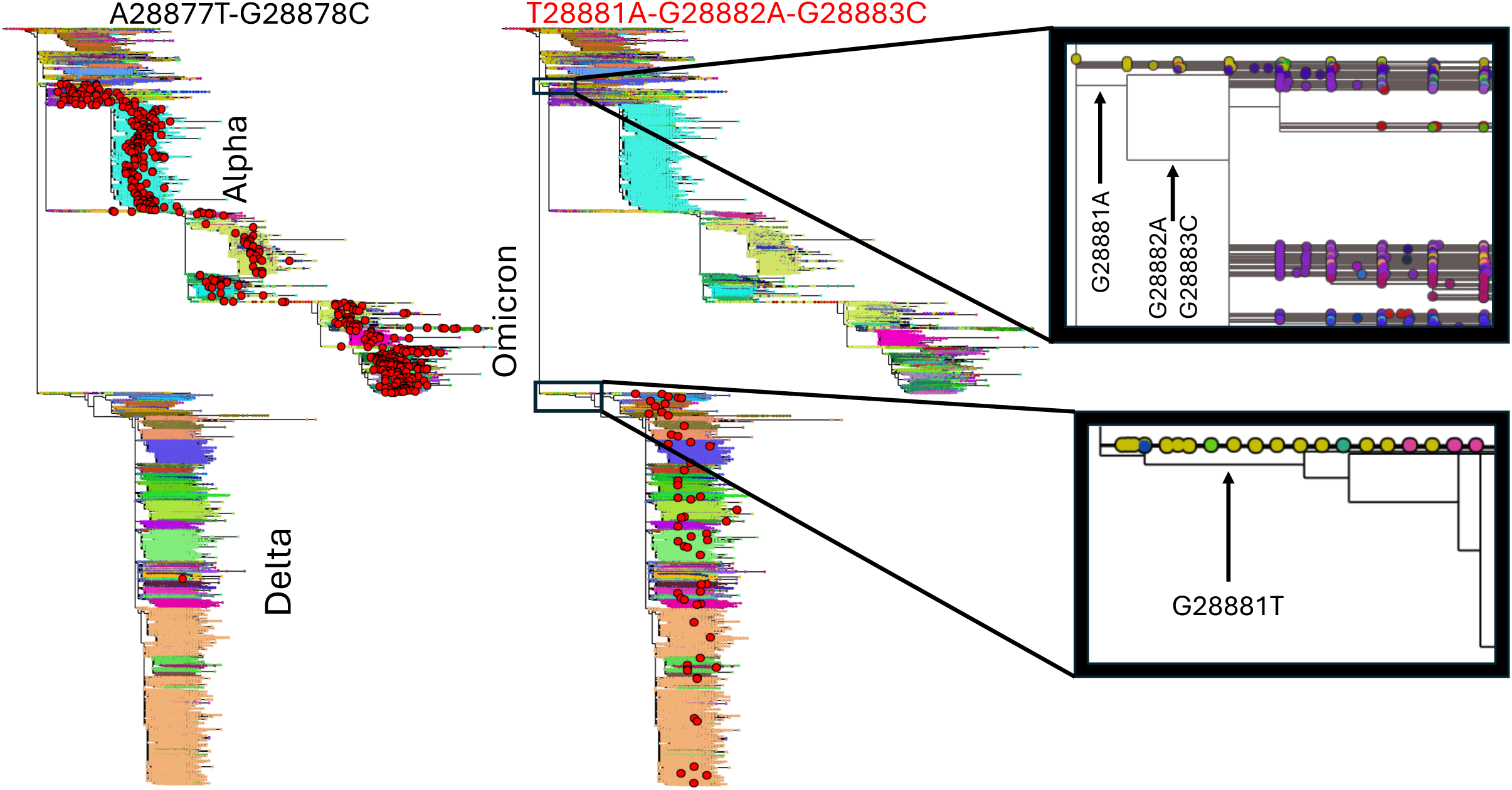
Phylogenetic distribution of MNSs A28877T-G28878C and T28881A-G28882A-G28883C. Details are as in Fig. 3. Left: occurrences of A28877T-G28878C are abundant in Alpha and Omicron lineages, but not in Delta. Right: occurrences of T28881A-G28882A-G28883C. While most occurrences of T28881A-G28882A-G28883C in Delta are likely to be artefacts (see section 2.2.2), we do not challenge the genuineness of the G28881A, G28882A and G28883C substitutions ancestral to the Alpha and Omicron lineages (top zoom-in box on the right), which cause most Alpha and Omicron genomes (in which most A28877T-G28878C MNSs occur) to have nucleotides AAC at positions 28881-28883.

### 2.3 Recombination and contamination

Homologous recombination can cause the recurrence of a MNS by transferring genetic material from a lineage with the MNS to a lineage without it (Fig. 1D). Such recombination-driven recurrences are expected to be more common for high-frequency MNSs (those that are present in many genomes), since recombination requires a host to be simultaneously infected by two viral lineages, one containing the MNS and one not containing it. Sample contamination can lead to a similar pattern (but only at the terminal branches of the phylogenetic tree, see Fig. 1D), causing genomes to appear recombinant, and is also expected to be more common for high-prevalence variants.

Unless these MNSs themselves would cause extremely high rates of recombination/contamination (which seems unlikely), then the rate of substitutions caused by recombination/contamination should depend only of the prevalence of the substitution, and substitutions with similar prevalence should have similar numbers of substitutions caused by recombi-nation/contamination. Because recurrent MNSs have similar prevalence to single-nucleotide substitutions in SARS-CoV-2 (Fig. 2C), we expect a similar number of recombination-induced substitutions for them as for single-nucleotide substitutions. However, recurrent MNSs show higher numbers of substitutions compared to single-nucleotide substitutions (Fig. 2C), and previous analyses suggest that recombination is not the leading cause of substitutions in SARS-CoV-2 (see e.g. [45]). It is therefore not plausible that recombination or contamination are the main driver of recurrent MNSs.

The only two recurrent MNSs with high prevalence are T28881A-G28882A-G28883C and G27382C-A27383T-T27384C. Of these, T28881A-G28882A-G28883C is in particular an ideal candidate for homologous recombination and contamination, given that it is present in close to half of the SARS-CoV-2 genomes. However, we only inferred 59 occurrences of T28881A-G28882A-G28883C, the vast majority of which appear to be artefacts caused by deletion masking (see section 2.2.2), not recombination or contamination. This further corroborates that it is unlikely that homologous recombination or contamination cause a substantial proportion of observed recurrent MNSs.

Assuming that homologous recombination and contamination events would not favor exchange of one genotype over the other, that is, they are symmetrical with respect to the parent genomes (note however that several factors might invalidate this assumption at certain positions, like context-dependent recombination rates or amplicon drop-out), one would expect them to cause a similar number of occurrences of a certain MNS as of its reverse. For example, considering A28881T-A28882G-C28883G as the reverse of T28881A-G28882A-G28883C (and also considering A28881G-A28882G-C28883G as its reverse, due to the high prevalence of nucleotide G at position 28881 (Fig. 4)), in total we inferred just 9 reversions of T28881A-G28882A-G28883C. In general, very few reversions were inferred for all recurrent MNSs (Table S2 column “Reversions”), further reducing the likelihood of a substantial impact from homologous recombination and contamination. For the other high-prevalence MNS, G27382C-A27383T-T27384C, we inferred only 5 reversions, in contrast with its 253 inferred occurrences. All these observations suggest that recombination and contamination are not the main cause of the recurrent MNSs considered.

### 2.4 Mechanisms of recurrent MNMs

Template switching is used by coronaviruses and most *Nidovirales* viruses during transcription to create a set of nested negative strand subgenomic RNAs (–sgRNAs)[46–48]. During template switching, negative strand synthesis stops at one of several transcription regulatory sequences (TRS-B, see Fig. 5A) and is re-initiated at the leader TRS (TRS-L) near the 5*^′^* end of the genome (Fig. 5B,C). After the transcription of the –sgRNA is completed (Fig. 5D), this is then used to synthesize a corresponding +sgRNA, which in turn is translated into a structural or accessory protein[46–48]. Template switch may be initiated at any of several TRS-Bs along the SARS-CoV-2 genome, resulting in –sgRNAs of different lengths, and this process is thought to modulate gene expression[46–48]. If template switching does not happen at any TRS-B, transcription results in a full genomic negative sense RNA (–gRNA), which is used as a template to synthesize +gRNA, which in turn can be used for translation or be packaged into new virions.

Template switching is also considered one of the main causes of indels, recombination, and non-canonical transcripts (uncommon –sgRNAs resulting from template switching from non-standard TRSs) in SARS-CoV-2[48–53], and it is therefore represents a reasonable potential mechanism causing recurrent MNMs.

**Figure 5:**
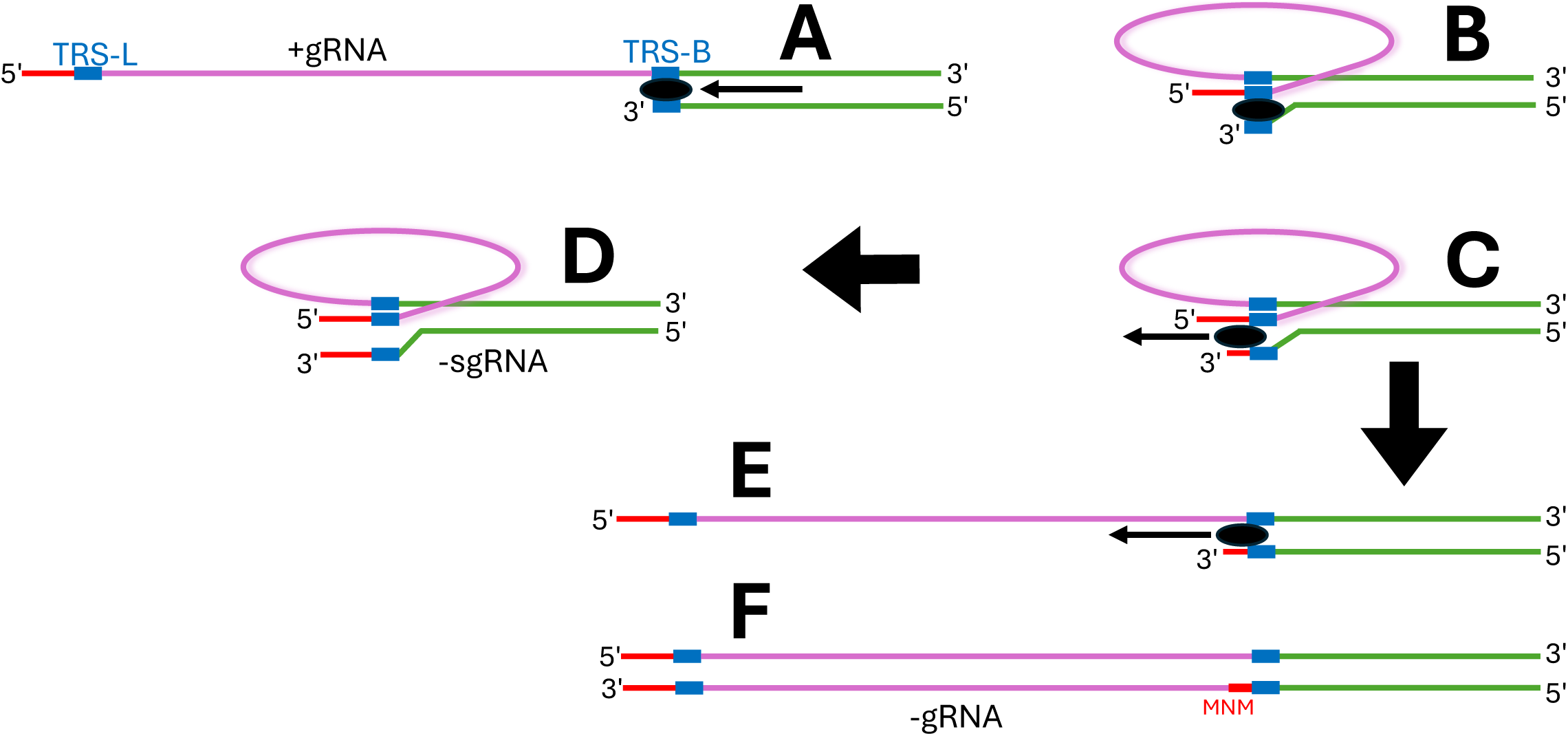
Interrupted template switching mechanism for recurrent MNMs. We sketch a possible mechanism for explaining the origin of most recurrent MNMs found in this study. **A** RNA-dependent RNA polymerase (RdRp, black oval) starts synthesizing a negative sense copy of the viral genome, and pauses at a TRS-B. **B** Template switching is initiated, RdRp is moved to the TRS-L, and **C** starts replicating nucleotides 5*^′^* of the TRS-L. After this, two scenarios can occur: either **D** template switching and replication are completed, resulting in a –sgRNA, or **E** template switching is interrupted, and RdRP returns to the original TRS-B and continues replication from there. **F** If after this template switching is not completed at any TRS-B, RdRp completes replication of a –gRNA containing an MNM just 5*^′^* of a TRS-B.

We find strong links between the majority of the highly recurrent MNMs, and the 5*^′^*-ACGAAC-3*^′^* TRS-L motif, suggesting a connection between many recurrent MNMs and template switching. More specifically, we find that recurrent MNMs seem to typically occur within just a few base pairs 5*^′^* of TRSs, and that the TRS-L region is a suitable template for almost every recurrent MNM. Following this evidence, We suggest that most highly recurrent MNMs in SARS-CoV-2 are caused by interrupted template switching during viral replication (see Fig. 5C,E). We suggest that in this process template switching induces a few bases from the TRS-L to be transcribed, after which the template switching is interrupted and transcription is restarted from near the original TRS-B. If template switching does not happen for the rest of the transcription process, this results in a –gRNA with an MNM just 5*^′^* of the TRS-B involved (Fig. 5F). The four most recurrent MNMs (C21302T-C21304A-G21305A, C21304A-G21305A, A28877T-G28878C and G27382C-A27383T-T27384C), and four other recurrent MNMs (T27875C-C27881T-G27882C-C27883T, C27881T-G27882C-C27883T, A27038T-T27039A-C27040A and A21550C-A21551T) fit this suggested mechanism well (Figs. 6 and 7). In particular, the highly recurrent MNM A28877T-G28878C almost exclusively happens in a genetic background containing the TRS motif ACGAAC nearby (Fig. 6, cf. Fig. 4), further confirming that TRSs are crucial to the most highly recurrent MNMs.

**Figure 6:**
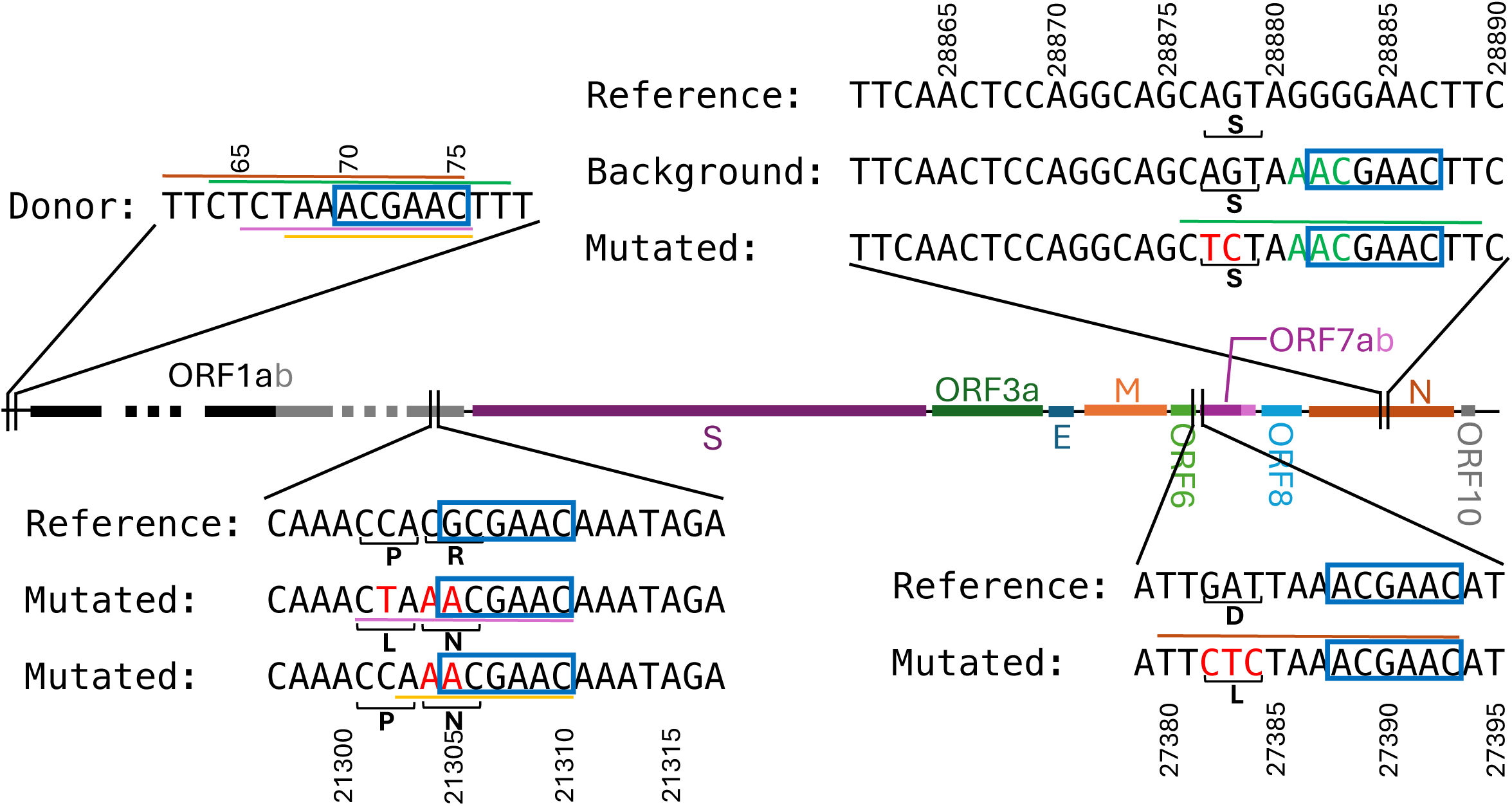
Suggested origin of the four most recurrent MNMs: C21302T-C21304A-G21305A, C21304A-G21305A, A28877T-G28878C and G27382C-A27383T-T27384C. TRS motifs are highlighted with blue boxes. Centre: Representation of the SARS-CoV-2 genome, annotated with its genes. Top left: 5*^′^* UTR reference genome sequence including the TRS-L (“Donor”). Top right: Representation of the A28877T-G28878C MNM, showing the reference sequence near the MNM, the background sequence that is typically mutated by the MNM (the differences between the reference and the background are highlighted in green), and the sequence resulting from the MNM (“Mutated”) with substituted nucleotides highlighted in red. Note that the background sequence contains the TRS motif, while the reference sequence does not (Fig. 4), which we suggest is the reason why the MNM mostly occurs in this background. Sequence identity between the mutated sequence and the TRS-L region is highlighted with a green line. Bottom right: Representation of the G27382C-A27383T-T27384C MNM. Sequence identity with the TRS-L region is highlighted with a brown line. Bottom left: Representation of the C21302T-C21304A-G21305A and C21304A-G21305A MNMs. Amino acids affected by the MNMs are shown under the nucleotide sequences. Sequence identity with the TRS-L region is highlighted with purple and yellow lines.

**Figure 7:**
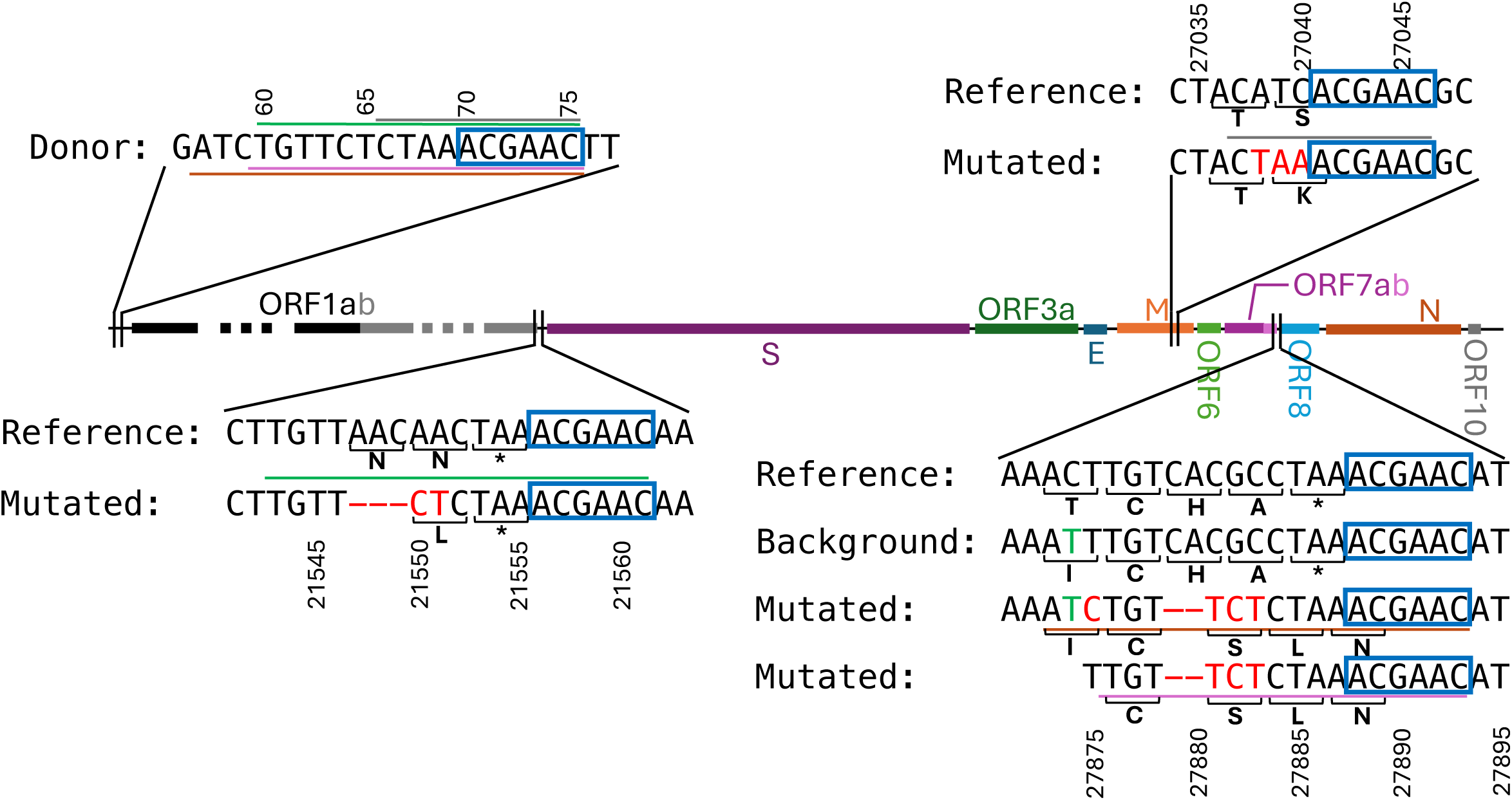
Suggested origin of MNMs A27038T-T27039A-C27040A, A21550C-A21551T, T27875C-C27881T-G27882C-C27883T and C27881T-G27882C-C27883T. Details are as in Fig. 6 The 2-nt deletions observed jointly with MNMs T27875C-C27881T-G27882C-C27883T and C27881T-G27882C-C27883T cause a frameshift in ORF7b that likely renders it non-functional.

Another six MNMs are consistent with the proposed mutational mechanism, although five of these MNMs occur at putative non-consensus TRSs (A<u>A</u>GAAC, ACGA<u>T</u>C, <u>G</u>CGAAC, <u>TT</u>GAAC, and AC<u>A</u>AAC, see Supplementary Figs. S4 and S5), while T26491C-A26492T-T26497C occurs at the 3*^′^* side of a TRS (Supplementary Fig. S4) instead of the usual 5*^′^* side. We cannot exclude that a different mechanism other than interrupted template switching might be the cause of some of these MNMs, and we do find, for some of them, other regions of sequence identity within the SARS-CoV-2 genome (Supplementary Fig. S6). This is particularly relevant for A20284T-T20285C, which is the only recurrent MNM considered here for which we found no obvious links to TRSs and template switching (Supplementary Fig. S6G).

### 2.5 Impact of recurrent MNMs on the inference of recombination

Just as homologous recombination can cause MNS occurrences (Fig. 1D), so recurrent MNMs might be misinterpreted as the result of homologous recombination events, since recurrent MNMs cause the same cluster of nearby substitutions to appear in different lineages. We typically model nucleotide substitutions as single-nucleotide independent events, and clustered substitutions within a small genomic region and within a phylogenetic branch are considered the hallmark of detectable homologous recombination events[45, 54]. This same pattern is however also caused by recurrent MNMs. A single recurrent MNM therefore might, on its own, generate sufficient signal to cause the spurious inference of many recombination events deemed statistically significant.

To test this, we investigated recombination events inferred by RIVET[39, 45] from global SARS-CoV-2 genomes collected up to June 2024[39]. Out of a total of 13,204 inferred recombination events, 1,858 (14.1%) were exclusively informed by one of the recurrent MNMs from Table 1. We therefore suggest that these are false positives, and that about one out of 7 inferred recombination events in SARS-CoV-2 evolution are likely the result of a recurrent MNM. In general, one would expect larger MNMs (those causing more simultaneous nucleotide substitutions) to generate more significant recombination event false positives. Indeed, since RIVET uses a parsimony score threshold of 3 to infer recombination events, 2-nt MNMs on their own do not lead to recombination event inference in RIVET, and did not contribute to any of the 1,858 recombination events informed exclusively by recurrent MNMs (Table 2).

**Table 2:** Number of recombination events inferred by RIVET involving recurrent MNSs. As in Table 1, we mark in red MNSs that are unlikely to be caused by recurrent MNMs. Recombinations involving MNS: numbers of recombination events containing the considered MNS within the recombinant genetic material. These are all the inferred recombinations that might have been influenced by recurrent MNSs. Recombinations entirely defined by MNS: number of inferred recombination events exclusively informed by the considered MNS (no other SNP is inferred to have been exchanged during recombination). We consider these events as recombination false positives caused by MNMs (except for the row corresponding to T28881A-G28882A-G28883C). Recombinations in this column are also included in the second column. 2-nt MNSs are not annotated at this column since RIVET uses a threshold of 3 substitutions for inferring recombination.

| MNS | Recombinations involving MNS | Recombinations entirely defined by MNS |
| --- | --- | --- |
| C21302T<br>-C21304A<br>-G21305A | 332 | 325 |
| C21304A<br>-G21305A | 6 |  |
| A28877T<br>-G28878C | 112 |  |
| G27382C<br>-A27383T<br>-T27384C | 850 | 575 |
| T26491C<br>-A26492T<br>-T26497C | 261 | 206 |
| G27758A<br>-T27760A | 0 |  |
| T27875C<br>-C27881T<br>-G27882C<br>-C27883T | 52 | 50 |
| C27881T<br>-G27882C<br>-C27883T | 41 | 16 |
| C25162A<br>-C25163A | 3 |  |
| T21294A<br>-G21295A<br>-G21296A | 320 | 287 |
| A27038T<br>-T27039A<br>-C27040A | 130 | 120 |
| A507T<br>-T508C<br>-G509A | 0 |  |
| T28881A<br>-G28882A<br>-G28883C | 542 | 279 |
| A21550C<br>-A21551T | 0 |  |
| T21994C<br>-T21995C | 9 |  |
| C13423A<br>-C13424A | 24 |  |
| A4576T<br>-T4579A | 11 |  |
| A20284T<br>-T20285C | 2 |  |
| G11083T<br>-C21575T | 0 |  |

Recurrent multi-nucleotide sequence errors, just like recurrent MNMs, might also lead to recombination inference false positives. Unlike the SARS-CoV-2 datasets we used, the genome sequences used by RIVET are not called with Viridian. For this reason, issues with reference biases mentioned earlier regarding T28881A-G28882A-G28883C might be more pronounced in the RIVET dataset, and are probably the cause of the large number of inferred recombination events for this MNS in Table 2.

These analyses show that recurrent MNMs and recurrent multi-nucleotide errors, if unaccounted for, greatly and adversely affect the inference of homologous recombination and cause high rates of false positives in SARS-CoV-2. However, recurrent MNMs and multi-nucleotide errors can also be flagged either by using Table 1, or, more generally, by classifying as unreliable all recombinations informed by clusters of substitutions *<* 10bp apart. We have now implemented both flags within RIVET v0.2.0 (https://github.com/TurakhiaLab/rivet).

## 3 Discussion

The existence and prevalence of MNMs has long been debated in the scientific literature[4, 5, 7, 8, 11–17]. MNMs allow evolution to explore areas of genome space that are less reachable through single-nucleotide substitutions alone[4], and substantially contribute to human disease[21, 22]. MNMs are also important in computational biology since they can negatively affect data analyses including annotation and diagnosis[21–25], and selection scans[26–29], since most methods wrongly interpret MNMs as combinations of separate single-nucleotide substitutions.

Previous studies have assumed that MNMs are rare and homogeneously distributed along the genome[27–30]. However, our analysis of millions of SARS-CoV-2 genomes shows the existence of several highly recurrent MNMs. The impact of highly recurrent MNMs in computational biology is considerably more significant than that of homogeneously distributed MNMs. For example, highly recurrent MNMs are expected to greatly compound the problem of false positives in positive selection scans, causing false positives not only with branch-site tests[55], but also with the generally more robust site-based tests[56].

As shown in section 2.5, highly recurrent MNMs also adversely affect the inference of recombination. Although our results are based on inference by RIVET[39, 45], we expect virtually all existing recombination detection methods to be affected by this issue.

Phylogenetic tree inference can also be negatively impacted by recurrent MNMs, since sequences descending from different occurrences of the same MNM could be wrongly clustered together and be interpreted as monophyletic due to the shared MNM. This is similar to the phylogenetic issues caused by convergent evolution[57], and might indeed be affecting our own analyses, leading to underestimation of the number of occurrences of recurrent MNMs.

We propose a mechanism of interrupted template switching as the leading cause of recurrent MNMs in SARS-CoV-2. This mechanism is consistent with the known process of transcription and replication of SARS-CoV-2[46–48], and might also explain the previously observed enrichment in indels near SARS-CoV-2 template switching hotspots[50, 51]. Since all coronaviruses (as well as other nidoviruses) share a similar template switching mechanism with SARS-CoV-2[58], it is fair to assume that their evolution is probably also affected by recurrent MNMs. We think it is an important open question to find which other organisms outside the *Nidovirales* also experience recurrent MNMs. As more genomic data is produced and shared for a large number of pathogens, we will not only be more able to address this question, but also be able to improve genomic analyses by accounting for identified recurrent MNMs.

While compensatory mutations[9] or positive selection[10] can also lead to MNSs, given the facts that SARS-CoV-2 phylogenetic tree branches are very short, that most highly recurrent MNMs we identified include synonymous or non-coding individual-nucleotide substitutions (Figs. 6 and 7, and Supplementary Figs. S4 and S5), and that many highly recurrent MNSs are composed of *>* 2 nucleotide substitutions, it is unlikely that positive or compensatory selection contribute substantially to our findings. It is also unlikely that the identified recurrent MNMs are themselves under strong positive selection (and therefore that positive selection might be causing their recurrence), given that they give rise to similar proportions of singletons (Fig. 2B) and descendants per mutation (Fig. 2C) as typical single-nucleotide mutations in the SARS-CoV-2 genome, and so appear to be under similar selective pressure.

The public availability of millions of SARS-CoV-2 genomes has made it possible to track the evolution of the virus in great detail during the COVID-19 pandemic. Without so many genomes, we would not have been able to identify highly recurrent MNMs, and without publicly available sequencing read data we would not have been able to validate them. In the future, we expect that further increases in sequencing capacity and genome data sharing will allow us to similarly track and understand the evolution of many other pathogens.

## 4 Acknowledgments

We thank Martin Hunt for help in investigating potential consensus sequence calling issues, and Yatish Turakhia for insightful discussions over the effects of MNMs on the inference of recombination events.

## 5 Methods

We investigated a SARS-CoV-2 dataset containing 2,072,111 genomes collected up to February 2023. Consensus sequences were called with Viridian[35], a tool that prevents common reference biases in genomic regions of low sequencing coverage. To reduce the number of contaminated samples, we removed from the dataset genomes enriched in within-sample genetic diversity, as described in [34]. We then masked alignment columns affected by previously identified recurrent sequence errors[34], and estimated a phylogenetic tree using MAPLE v0.6.8 with an UNREST substitution model, rate variation, and deep SPR phylogenetic search. For a full description of data preparation and phylogenetic inference, see [34]. We then kept track of all substitutions inferred along the phylogenetic tree by MAPLE with posterior probability above 95%.

We counted the number of occurrences of single-nucleotide substitutions and their number of descendants in the tree. We then did the same calculations for all pairs of single-nucleotide substitutions found on the same branch in the tree, even those occurring in different parts of the genome. Due to the short branches in the SARS-CoV-2 phylogenetic tree, a manageable number of such pairs was observed. When recurrent clusters of *>* 2 nucleotide substitutions where found, these were also tracked (and occurrences of larger clusters where not counted towards the numbers of occurrences of smaller nested clusters).

We identified all genomes descending from any occurrence of the most recurrent MNSs (those in Table 1) on a terminal branch of the tree. For any of these genomes that were part of a cherry (a 2-tip subtree) in the phylogenetic tree, we analyzed sequencing read data (as represented in Viridian quality control files) for both members of the cherry. We wrote a custom python script to extract information about coverage, indels, and variants near the considered MNSs in these genomes.

The phylogenetic distribution of recurrent MNSs was visually investigated using Taxonium v2.0.115[44].

The output of RIVET was analyzed with a custom python script to extract putative recombination events informed by the MNSs in Table 1.

The alignment, metadata, and inferred tree are publicly available on Zenodo[59]. The list of recombination events inferred by RIVET, and the list of cherry samples analyzed here, are available from https://github.com/NicolaDM/MAPLE/tree/main/multinucleotideMutations.

## 6 Supplementary Figures

**Figure S1:**
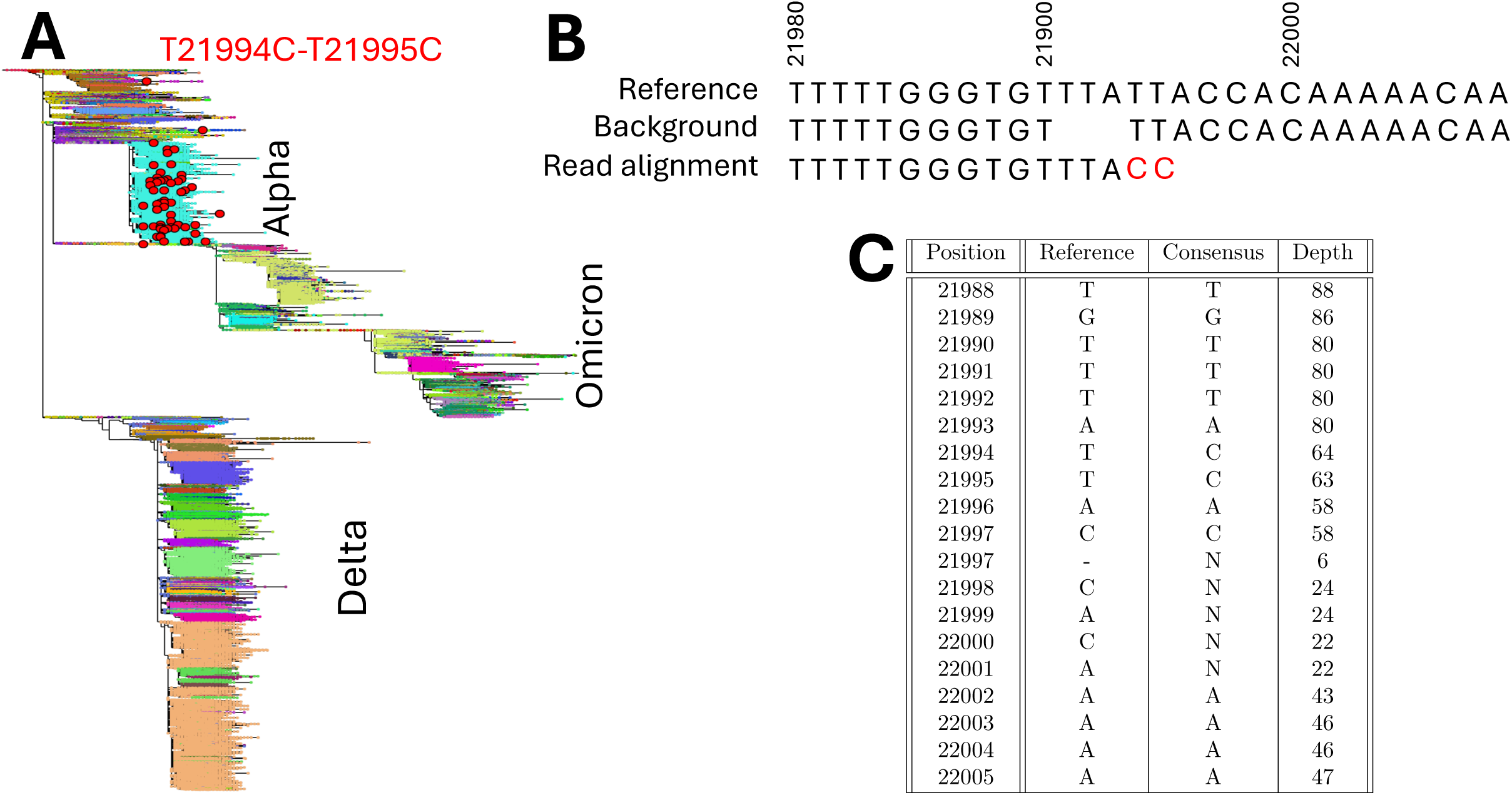
Issues with MNS T21994C-T21995C. **A** SARS-COV-2 phylogenetic tree annotated (red circles) with occurrences of T21994C-T21995C MNSs. Details are as in Fig. 3. This MNS almost exclusively occurs in the Alpha lineage, where a deletion at positions 21991–21993 is ancestral to most genomes. **B** Local reference genome sequence (“Reference”), the background genome in Alpha (“Background”, showing the deletion at positions 21991–21993 prevalent in the Alpha variant), and an example of a read alignment observed in samples containing the T21994C-T21995C MNS (“Read alignment”). The mis-aligned nucleotides causing the T21994C-T21995C MNS in the consensus genome are highlighted in red. These are typically found at the 3*^′^* end of the read, where read alignment can favor 2 substitutions over a 3-nt deletion. **C** Viridian[35] quality control data for a random sample (ERR6719906) among the 28 containing a singleton T21994C-T21995C MNS and that are part of a phylogenetic cherry. We show reference position, reference nucleotide, Viridian consensus nucleotide, and total depth (a subset of the fields shown in Supplementary Table S1). Read alignment errors shift read depth to the deleted positions 21991–21993, making them appear not deleted in the consensus sequence, causing the T21994C-T21995C MNS, and leaving insufficient depth at positions 21998–22001 to call the reference genome. While this that represents only one example genome, Supplementary Table S2 shows that 26/28 considered samples descending from a singleton T21994C-T21995C MNS and part of a cherry present low depth positions near the MNS, and visual inspection confirms that most of them have a similar pattern in read data alignment to that shown here.

**Figure S2:**
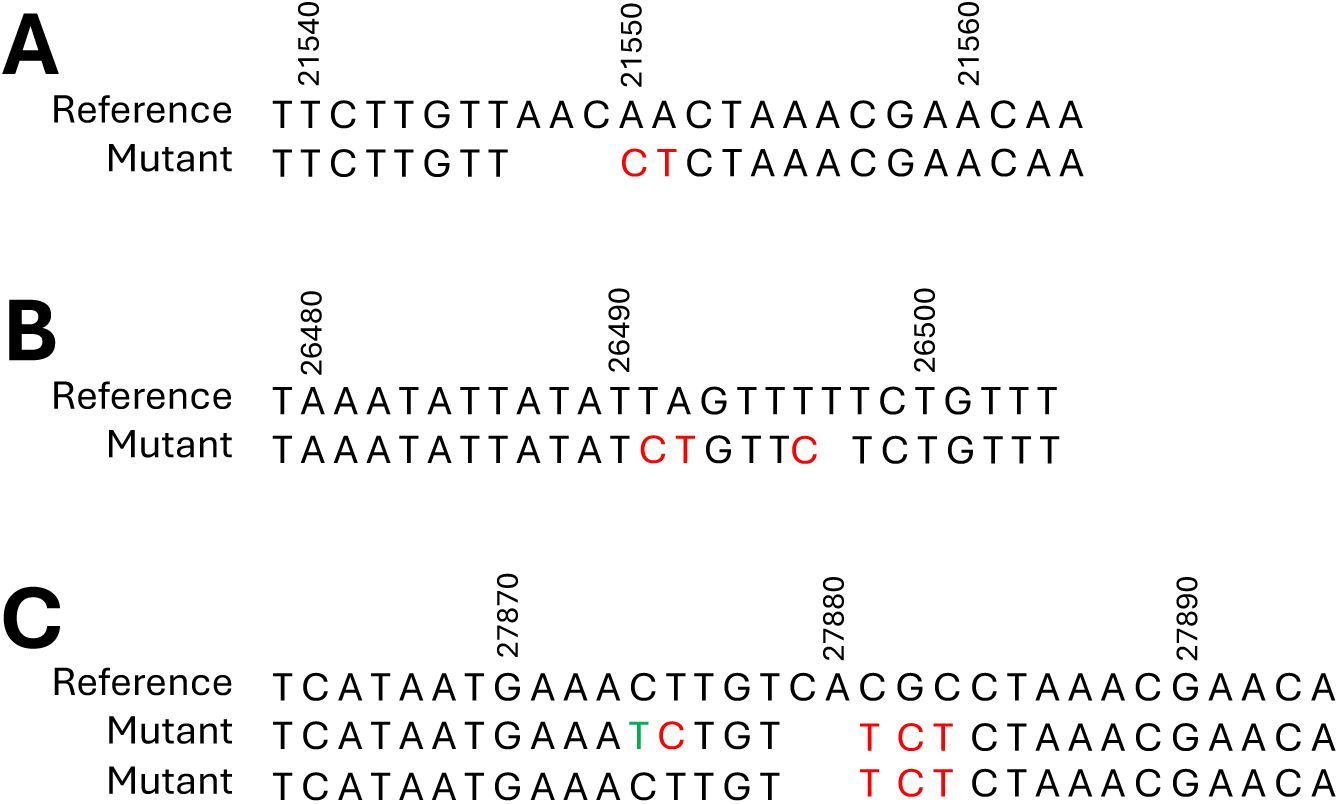
Complex MNMs. Some recurrent MNMs appear to involve deletions as well as substitutions. We show the reference genome sequence (“Reference”) near the MNMs considered, as well as the sequence mutated by the MNM (“Mutant”). Nucleotide substitutions defining the MNM are marked in red. A MNM A21550C-A21551T involves a 3-nucleotide deletion. B MNM T26491C-A26492T-T26497C involves a 1-nucleotide deletion. C MNMs T27875C-C27881T-G27882C-C27883T and C27881T-G27882C-C27883T involve 2-nucleotide deletions; the green T in the T27875C-C27881T-G27882C-C27883T mutated sequence is part of the genomic background where T27875C-C27881T-G27882C-C27883T occurs (see Fig. 3).

**Figure S3:**
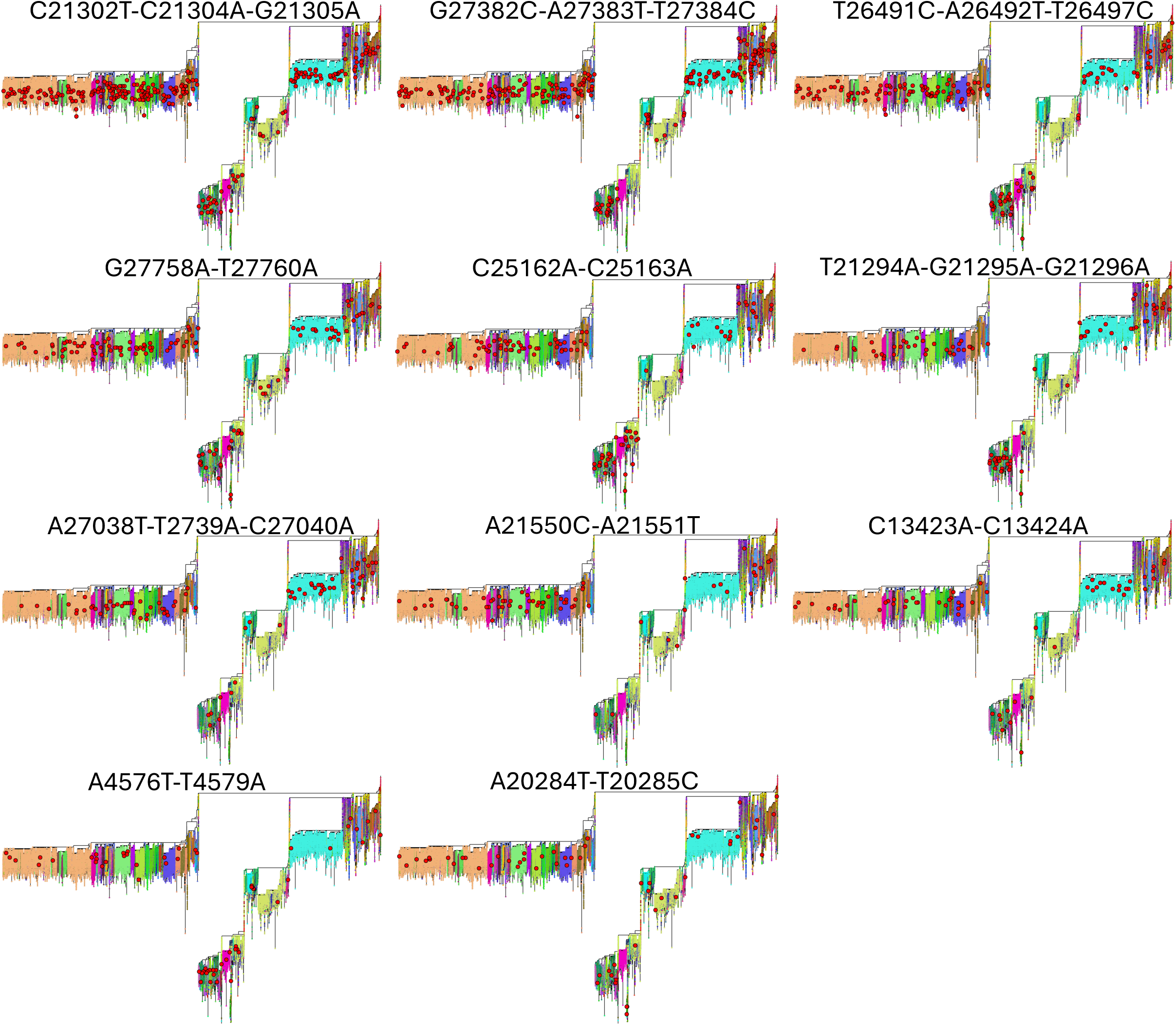
Phylogenetic distributions of recurrent MNSs. Each copy of the SARS-CoV-2 phylogenetic tree (details as in Fig. 3) is annotated (red circles) with occurrences of a different recurrent MNS. These MNSs do not show obvious phylogenetic clustering.

**Figure S4:**
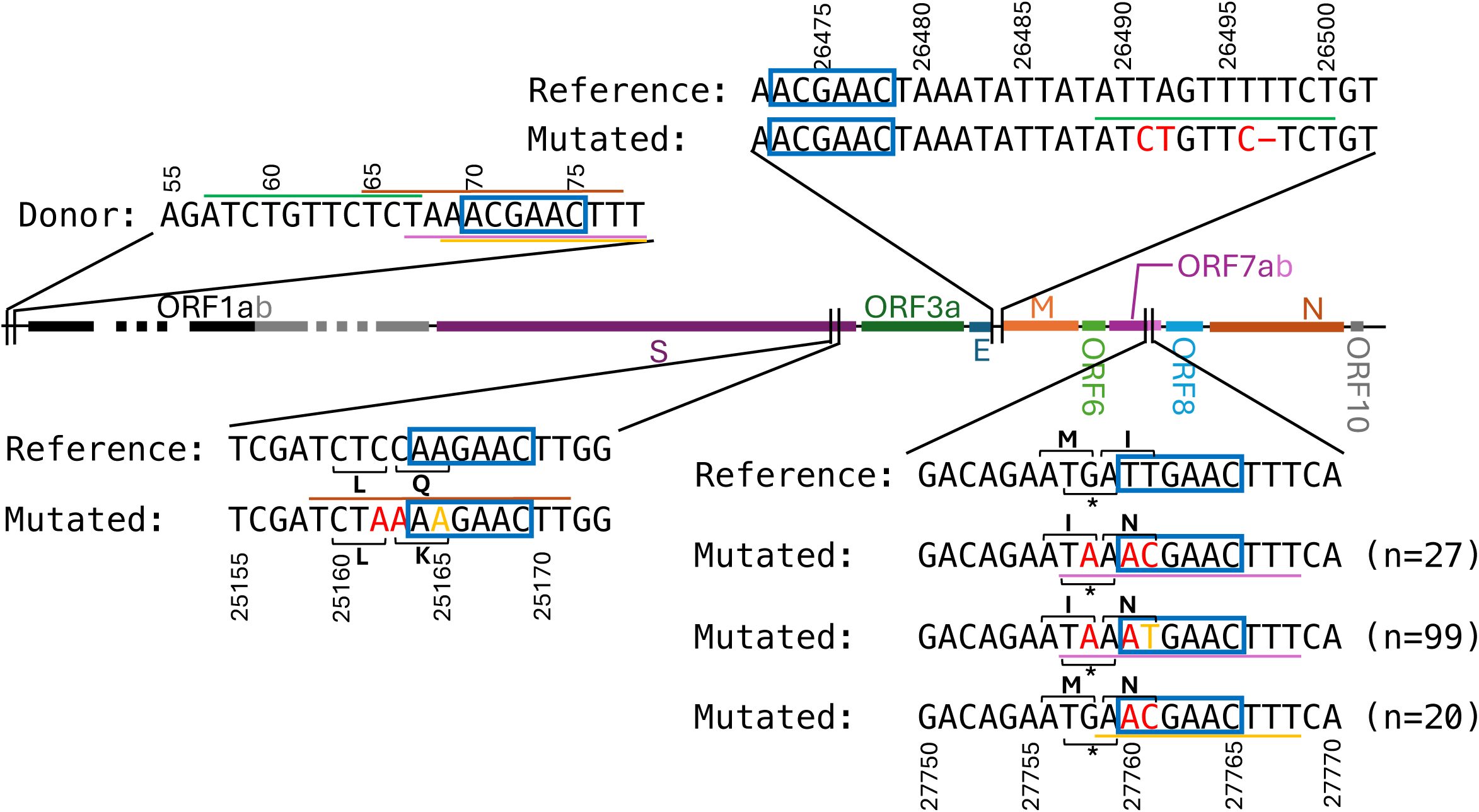
Suggested origin of MNMs T26491C-A26492T-T26497C, G27758A-T27760A, and C25162A-C25163A. Details are as in Fig. 6. Orange nucleotides are mismatches with respect to the highlighted TRS-L region of identity. At bottom right, we show G27758A-T27760A together with its extended version G27758A-T27760A-T27761C (27 occurrences out of all 126 occurrences of G27758A-T27760A) and its alternative T27760A-T27761C (20 occurrences) which are not recurrent enough to be listed in Table 1 on their own. These sequences are annotated with their translations as the end of ORF7a (below sequence) and the start of ORF7b (above). The M1I amino acid substitution caused by G27758A appears likely to entirely remove ORF7b.

**Figure S5:**
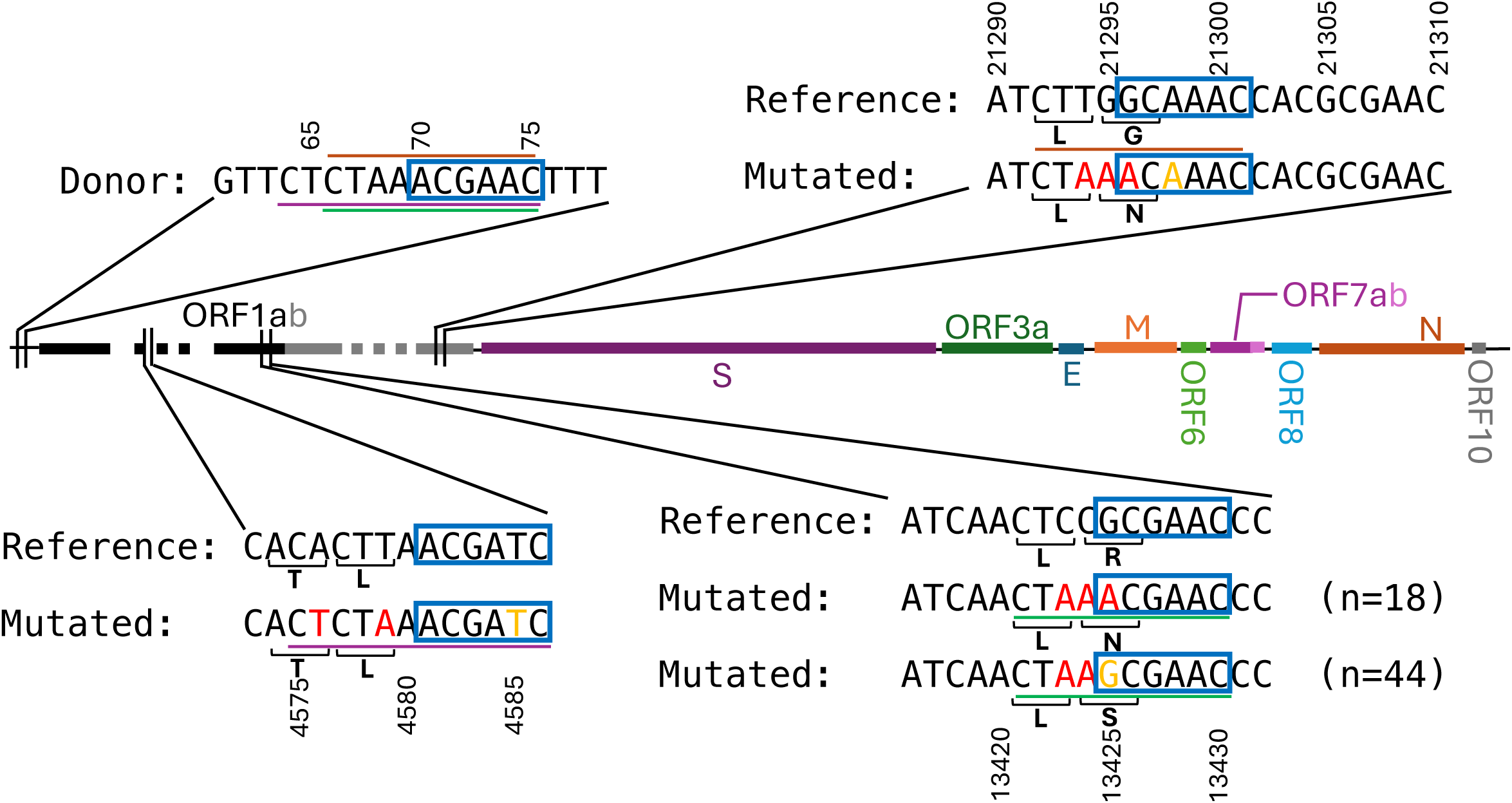
Suggested origin of MNMs T21294A-G21295A-G21296A, C13423A-C13424A and A4576T-T4579A. Details are as in Fig. 6. Orange nucleotides are mismatches with respect to the highlighted TRS-L region of identity. We show two versions of C13423A-C13424A, one of which includes also substitution G13425A and which is not recurrent enough (18 occurrences out of all 62 occurrences of the cluster) to be listed in Table 1 on its own.

**Figure S6:**
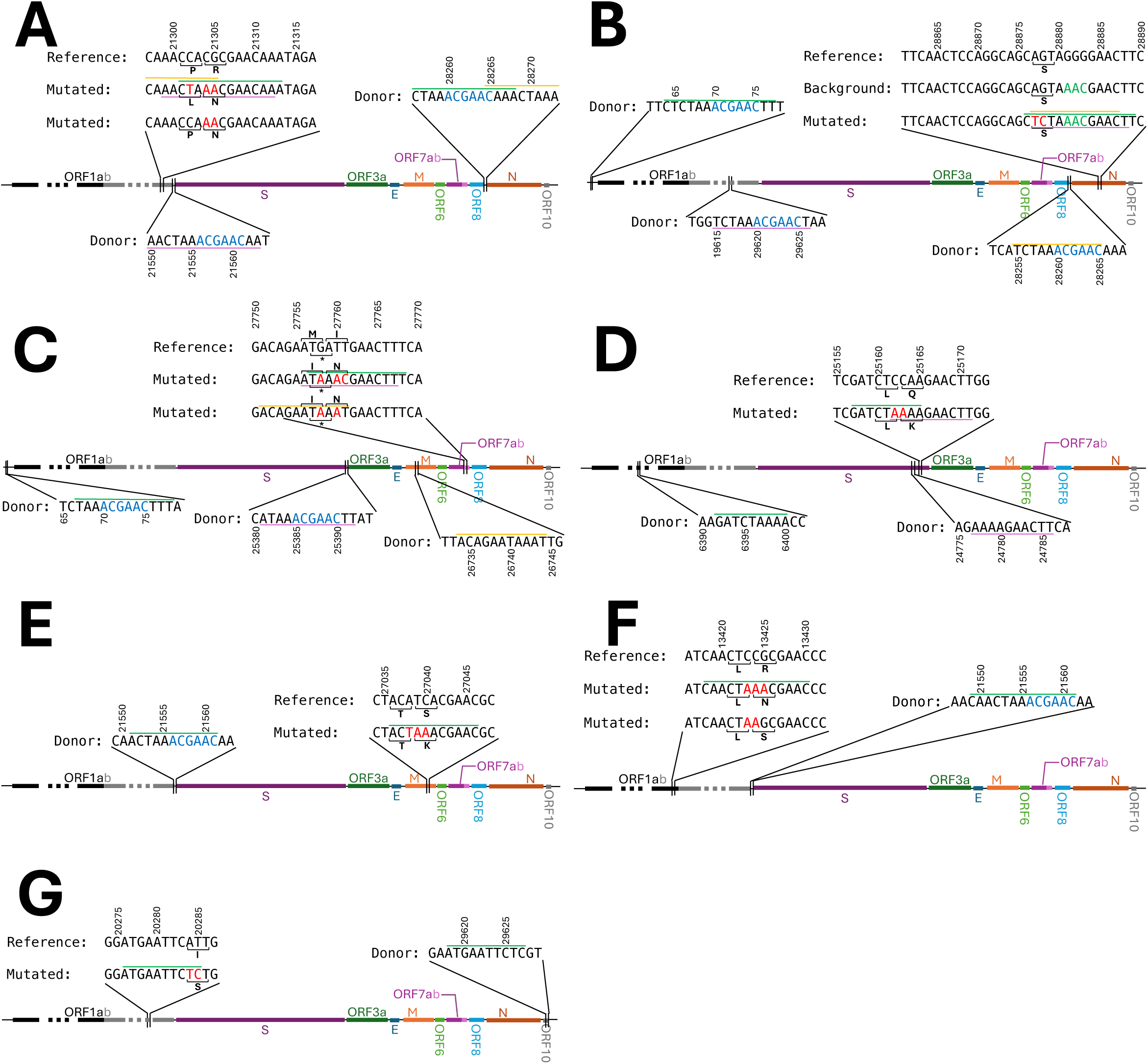
Possible other templates in the SARS-CoV-2 genome. Here we show regions of the SARS-CoV-2 genome, distinct from the TRS-L, that have long sequence identity (*≥* 9bp) with the mutated version of some of the recurrent MNMs. Details are as in Fig. 6. We highlight in blue the TRS consensus motif ACGAAC found in many of these regions. A C21302T-C21304A-G21305A and C21304A-G21305A (see also Fig. 6). B A28877T-G28878C (see also Fig. 6). C G27758A-T27760A (see also Supplementary Fig. S4). D C25162A-C25163A (see also Supplementary Fig. S4). E A27038T-T27039A-C27040A (see also Fig. 7). F C13423A-C13424A (see also Supplementary Fig. S5). G A20284T-T20285C, the only recurrent MNM we could not explain through interrupted template switching.

## 7 Supplementary Tables

**Table S1:** Issues in sequencing read data for sample SRR19943422. We show the Viridian[35] quality control data for a random sample among the 32 containing a singleton A507T-T508C-G509A MNS and that are part of a phylogenetic cherry. We show only data for positions near the considered MNS. Position: reference genome position. Reference: reference nucleotide at the position. Consensus: consensus nucleotide called by Viridian from the read data at the considered position for the considered sample. Depth: sequencing read depth at the position. Clean depth: depth after trimming primers from the reads. Consensus clean depth: clean depth supporting the Viridian consensus nucleotide at the position. Amplicons: amplicons covering the position for the sequencing protocol inferred by Viridian. Primers: primers covering the position for the sequencing scheme inferred by Viridian. The table shows that following a region of high depth up to reference position 506, depth suddenly drops, and read alignments contain deletions (unlike the sibling genomes in the phylogenetic cherry, not shown). Also, within-sample heterozygosity is observed at positions 504 and 505 (the consensus clean depth is substantially lower than the total clean depth). After position 536 the consensus sequence is masked by Viridian (the consensus nucleotide is N) due to primer r1_1.1.592753_r_0, and apparently due to dropout of amplicon r1_1.1.409836. All these observations suggest that the herozygosity and indels might be due to inaccurate primer identification and consequentially incomplete primer trimming. Aligning the consensus Viridian genome to the reference with MAFFT[60] (see details in [34]) introduces substitutions A507T, T508C, and G509A, and a deletion. While this table highlights just one example, Supplementary Table S2 shows that all considered samples descending from a singleton A507T-T508C-G509A MNS and part of a cherry also present heterozygosity and indels at these positions, and visual inspection of their read data further led us to conclude that this recurrent MNS is the consequence of sequence artefacts.

| Position | Reference | Consensus | Depth | Clean depth | Consensus clean depth | Amplicons | Primers |
| --- | --- | --- | --- | --- | --- | --- | --- |
| 497 | A | A | 1789 | 1279 | 1275 | r1_1.1.409836;r1_1.1.592753;r1_1.1.876634 | r1_1.1.876634_r_0;r1_1.1.409836_l_0 |
| 498 | C | C | 1789 | 1279 | 1271 | r1_1.1.409836;r1_1.1.592753;r1_1.1.876634 | r1_1.1.876634_r_0;r1_1.1.409836_l_0 |
| 499 | T | T | 1787 | 1278 | 1278 | r1_1.1.409836;r1_1.1.592753;r1_1.1.876634 | r1_1.1.876634_r_0;r1_1.1.409836_l_0 |
| 500 | G | G | 1787 | 1278 | 1276 | r1_1.1.409836;r1_1.1.592753;r1_1.1.876634 | r1_1.1.876634_r_0;r1_1.1.409836_l_0 |
| 501 | C | C | 1786 | 1277 | 1277 | r1_1.1.409836;r1_1.1.592753;r1_1.1.876634 | r1_1.1.876634_r_0;r1_1.1.409836_l_0 |
| 502 | A | A | 1785 | 1276 | 1275 | r1_1.1.409836;r1_1.1.592753;r1_1.1.876634 | r1_1.1.876634_r_0;r1_1.1.409836_l_0 |
| 503 | C | C | 1535 | 1028 | 1027 | r1_1.1.409836;r1_1.1.592753;r1_1.1.876634 | r1_1.1.876634_r_0;r1_1.1.409836_l_0 |
| 504 | C | C | 1751 | 1253 | 1004 | r1_1.1.409836;r1_1.1.592753;r1_1.1.876634 | r1_1.1.876634_r_0;r1_1.1.409836_l_0 |
| 505 | T | T | 1746 | 1249 | 999 | r1_1.1.409836;r1_1.1.592753;r1_1.1.876634 | r1_1.1.876634_r_0;r1_1.1.409836_l_0 |
| 506 | C | C | 1740 | 1246 | 1244 | r1_1.1.409836;r1_1.1.592753;r1_1.1.876634 | r1_1.1.876634_r_0;r1_1.1.409836_l_0 |
| 507 | A | - | . | . | . | r1_1.1.409836;r1_1.1.592753;r1_1.1.876634 | r1_1.1.876634_r_0;r1_1.1.409836_l_0 |
| 508 | T | - | . | . | . | r1_1.1.409836;r1_1.1.592753;r1_1.1.876634 | r1_1.1.876634_r_0;r1_1.1.409836_l_0 |
| 509 | G | - | . | . | . | r1_1.1.409836;r1_1.1.592753 | r1_1.1.409836_l_0 |
| 510 | G | - | . | . | . | r1_1.1.409836;r1_1.1.592753 | r1_1.1.409836_l_0 |
| 511 | T | T | 255 | 255 | 252 | r1_1.1.409836;r1_1.1.592753 | r1_1.1.409836_l_0 |
| 512 | C | C | 256 | 256 | 253 | r1_1.1.409836;r1_1.1.592753 | r1_1.1.409836_l_0 |
| 513 | A | A | 255 | 255 | 252 | r1_1.1.409836;r1_1.1.592753 | r1_1.1.409836_l_0 |
| 514 | T | - | . | . | . | r1_1.1.409836;r1_1.1.592753 | . |
| 515 | G | - | . | . | . | r1_1.1.409836;r1_1.1.592753 | . |
| 516 | T | - | . | . | . | r1_1.1.409836;r1_1.1.592753 | . |
| 517 | T | - | . | . | . | r1_1.1.409836;r1_1.1.592753 | . |
| 518 | A | - | . | . | . | r1_1.1.409836;r1_1.1.592753 | . |
| 519 | T | - | . | . | . | r1_1.1.409836;r1_1.1.592753 | . |
| 520 | G | - | . | . | . | r1_1.1.409836;r1_1.1.592753 | . |
| 521 | G | - | . | . | . | r1_1.1.409836;r1_1.1.592753 | . |
| 522 | T | - | . | . | . | r1_1.1.409836;r1_1.1.592753 | . |
| 523 | T | T | 253 | 253 | 252 | r1_1.1.409836;r1_1.1.592753 | . |
| 524 | G | G | 253 | 253 | 249 | r1_1.1.409836;r1_1.1.592753 | . |
| 525 | A | A | 253 | 253 | 252 | r1_1.1.409836;r1_1.1.592753 | . |
| 526 | G | G | 251 | 251 | 243 | r1_1.1.409836;r1_1.1.592753 | . |
| 527 | C | C | 253 | 253 | 250 | r1_1.1.409836;r1_1.1.592753 | . |
| 528 | T | T | 253 | 253 | 250 | r1_1.1.409836;r1_1.1.592753 | . |
| 529 | G | G | 253 | 253 | 253 | r1_1.1.409836;r1_1.1.592753 | . |
| 530 | G | G | 253 | 253 | 253 | r1_1.1.409836;r1_1.1.592753 | . |
| 531 | T | T | 253 | 253 | 252 | r1_1.1.409836;r1_1.1.592753 | . |
| 532 | A | A | 253 | 253 | 253 | r1_1.1.409836;r1_1.1.592753 | . |
| 533 | G | G | 253 | 253 | 253 | r1_1.1.409836;r1_1.1.592753 | . |
| 534 | C | C | 253 | 253 | 253 | r1_1.1.409836;r1_1.1.592753 | . |
| 535 | A | A | 252 | 252 | 252 | r1_1.1.409836;r1_1.1.592753 | . |
| 536 | G | N | 252 | 0 | 0 | r1_1.1.409836;r1_1.1.592753 | r1_1.1.592753_r_0 |
| 537 | A | N | 245 | 0 | 0 | r1_1.1.409836;r1_1.1.592753 | r1_1.1.592753_r_0 |
| 538 | A | N | 244 | 0 | 0 | r1_1.1.409836;r1_1.1.592753 | r1_1.1.592753_r_0 |
| 539 | C | N | 245 | 0 | 0 | r1_1.1.409836;r1_1.1.592753 | r1_1.1.592753_r_0 |
| 540 | T | N | 245 | 0 | 0 | r1_1.1.409836;r1_1.1.592753 | r1_1.1.592753_r_0 |
| 541 | C | N | 245 | 0 | 0 | r1_1.1.409836;r1_1.1.592753 | r1_1.1.592753_r_0 |
| 542 | G | N | 245 | 0 | 0 | r1_1.1.409836;r1_1.1.592753 | r1_1.1.592753_r_0 |
| 543 | A | N | 245 | 0 | 0 | r1_1.1.409836;r1_1.1.592753 | r1_1.1.592753_r_0 |
| 544 | A | N | 245 | 0 | 0 | r1_1.1.409836;r1_1.1.592753 | r1_1.1.592753_r_0 |
| 545 | G | N | 244 | 0 | 0 | r1_1.1.409836;r1_1.1.592753 | r1_1.1.592753_r_0 |

**Table S2:** Other features of highly recurrent MNSs. MNSs unlikely to be caused by recurrent MNMs are highlighted in red. Phylogenetic clustering: MNSs clearly non-homogeneously spread along the phylogenetic tree (as in Supplementary Fig. S3) are marked with X. Reversions: Number of flipped substitutions inferred for the considered MNS; e.g. A21304C-A21305G is the reversion of C21304A-G21305A. The last three columns list how many of the investigated samples containing singletons for the considered MNS also contain near the same MNS: deletions in the consensus sequence; low (*<* 100) read coverage positions; heterozygosities (*>* 5% non-consensus nucleotide frequency in the read alignment). We consider positions *≤*20bp before the first base in the MNS, or *≤*25bp after it. For example, for the first row, out of 78 samples containing a singleton C21302T-C21304A-G21305A and with a sibling sample in the tree, 5 contained low read coverage in at least one site between genome positions 21282 and 21327.

| MNS | Phylogenetic clustering | Reversions | Deletions in cherries | Low coverage in cherries | Heterozygosities in cherries |
| --- | --- | --- | --- | --- | --- |
| C21302T<br>-C21304A<br>-G21305A |  | 0 | 0/78 | 5/78 | 12/78 |
| C21304A<br>-G21305A |  | 0 | 0/97 | 4/97 | 6/97 |
| A28877T<br>-G28878C | X | 13 | 1/131 | 2/131 | 13/131 |
| G27382C<br>-A27383T<br>-T27384C |  | 5 | 0/69 | 1/69 | 5/69 |
| T26491C<br>-A26492T<br>-T26497C |  | 0 | 51/51 | 0/51 | 7/51 |
| G27758A<br>-T27760A |  | 0 | 0/37 | 1/37 | 6/37 |
| T27875C<br>-C27881T<br>-G27882C<br>-C27883T | X | 0 | 9/9 | 0/9 | 0/9 |
| C27881T<br>-G27882C<br>-C27883T |  | 0 | 23/23 | 0/23 | 4/23 |
| C25162A<br>-C25163A |  | 0 | 0/29 | 0/29 | 3/29 |
| T21294A<br>-G21295A<br>-G21296A |  | 0 | 0/21 | 1/21 | 0/21 |
| A27038T<br>-T27039A<br>-C27040A |  | 0 | 0/26 | 3/26 | 6/26 |
| A507T<br>-T508C<br>-G509A |  | 0 | 32/32 | 2/32 | 32/32 |
| T28881A<br>-G28882A<br>-G28883C |  | 9 | 12/16 | 0/16 | 3/16 |
| A21550C<br>-A21551T |  | 0 | 16/16 | 0/16 | 3/16 |
| T21994C<br>-T21995C | X | 2 | 7/28 | 26/28 | 0/28 |
| C13423A<br>-C13424A |  | 0 | 0/24 | 0/24 | 0/24 |
| A4576T<br>-T4579A |  | 0 | 0/14 | 4/14 | 0/14 |
| A20284T<br>-T20285C |  | 0 | 0/14 | 0/14 | 0/14 |
| G11083T<br>-C21575T |  | 0 | 0/14 | 0/14 | 14/14 |

## References

1. Kloosterman, W. P. et al. Characteristics of de novo structural changes in the human genome. Genome Research 25, 792–801 (2015).

2. Hamilton, M. B. Population Genetics (John Wiley & Sons, 2021).

3. Yang, Z. & Rannala, B. Molecular phylogenetics: principles and practice. Nature Reviews Genetics 13, 303–314 (2012).

4. Averof, M., Rokas, A., Wolfe, K. H. & Sharp, P. M. Evidence for a high frequency of simultaneous double-nucleotide substitutions. Science 287, 1283–1286 (2000).

5. Smith, N. G., Webster, M. T. & Ellegren, H. A low rate of simultaneous double-nucleotide mutations in primates. Molecular Biology and Evolution 20, 47–53 (2003).

6. Kosiol, C., Holmes, I. & Goldman, N. An empirical codon model for protein sequence evolution. Molecular Biology and Evolution 24, 1464–1479 (2007).

7. Golding, G. B. & Glickman, B. W. Sequence-directed mutagenesis: evidence from a phylogenetic history of human alpha-interferon genes. Proceedings of the National Academy of Sciences USA 82, 8577–8581 (1985).

8. Whelan, S. & Goldman, N. Estimating the frequency of events that cause multiple-nucleotide changes. Genetics 167, 2027–2043 (2004).

9. Belinky, F., Sela, I., Rogozin, I. B. & Koonin, E. V. Crossing fitness valleys via double substitutions within codons. BMC Biology 17, 1–15 (2019).

10. Bazykin, G. A., Kondrashov, F. A., Ogurtsov, A. Y., Sunyaev, S. & Kondrashov, A. S. Positive selection at sites of multiple amino acid replacements since rat–mouse divergence. Nature 429, 558–562 (2004).

11. Hampsey, D. M., Ernst, J. F., Stewart, J. W. & Sherman, F. Multiple base-pair mutations in yeast. Journal of Molecular Biology 201, 471– 486 (1988).

12. Nakazawa, H. et al. UV and skin cancer: specific p53 gene mutation in normal skin as a biologically relevant exposure measurement. Proceedings of the National Academy of Sciences USA 91, 360–364 (1994).

13. Hodgkinson, A. & Eyre-Walker, A. Human triallelic sites: evidence for a new mutational mechanism? Genetics 184, 233–241 (2010).

14. Schrider, D. R., Hourmozdi, J. N. & Hahn, M. W. Pervasive multinucleotide mutational events in eukaryotes. Current Biology 21, 1051– 1054 (2011).

15. Chen, J.-M., Cooper, D. N. & Férec, C. A new and more accurate estimate of the rate of concurrent tandem-base substitution mutations in the human germline: 0.4% of the single-nucleotide substitution mutation rate. Human Mutation 35, 392–394 (2014).

16. Assaf, Z. J., Tilk, S., Park, J., Siegal, M. L. & Petrov, D. A. Deep sequencing of natural and experimental populations of *Drosophila melanogaster* reveals biases in the spectrum of new mutations. Genome Research 27, 1988–2000 (2017).

17. Wang, Q. et al. Landscape of multi-nucleotide variants in 125,748 human exomes and 15,708 genomes. Nature Communications 11, 2539 (2020).

18. Löytynoja, A. & Goldman, N. Short template switch events explain mutation clusters in the human genome. Genome Research 27, 1039– 1049 (2017).

19. Walker, C. R., Scally, A., De Maio, N. & Goldman, N. Short-range template switching in great ape genomes explored using pair hidden Markov models. PLoS Genetics 17, e1009221 (2021).

20. Löytynoja, A. Thousands of human mutation clusters are explained by short-range template switching. Genome Research 32, 1437–1447 (2022).

21. Kaplanis, J. et al. Exome-wide assessment of the functional impact and pathogenicity of multinucleotide mutations. Genome Research 29, 1047–1056 (2019).

22. Wang, D. et al. Pancan-MNVQTLdb: systematic identification of multi-nucleotide variant quantitative trait loci in 33 cancer types. NAR Cancer 4, zcac043 (2022).

23. Degalez, F. et al. Watch out for a second SNP: focus on multi-nucleotide variants in coding regions and rescued stop-gained. Frontiers in Genetics 12, 659287 (2021).

24. Wakeling, M. N. et al. Misannotation of multiple-nucleotide variants risks misdiagnosis. Wellcome Open Research 4 (2019).

25. Srinivasan, S. et al. Misannotated multi-nucleotide variants in public cancer genomics datasets lead to inaccurate mutation calls with significant implications. Cancer Research 81, 282–288 (2021).

26. Venkat, A., Hahn, M. W. & Thornton, J. W. Multinucleotide mutations cause false inferences of lineage-specific positive selection. Nature Ecology & Evolution 2, 1280–1288 (2018).

27. Dunn, K. A., Kenney, T., Gu, H. & Bielawski, J. P. Improved inference of site-specific positive selection under a generalized parametric codon model when there are multinucleotide mutations and multiple nonsynonymous rates. BMC Evolutionary Biology 19, 1–19 (2019).

28. Lucaci, A. G., Wisotsky, S. R., Shank, S. D., Weaver, S. & Kosakovsky Pond, S. L. Extra base hits: widespread empirical support for instantaneous multiple-nucleotide changes. PloS One 16, e0248337 (2021).

29. Lucaci, A. G., Zehr, J. D., Enard, D., Thornton, J. W. & Kosakovsky Pond, S. L. Evolutionary shortcuts via multinucleotide substitutions and their impact on natural selection analyses. Molecular Biology and Evolution 40, msad150 (2023).

30. De Maio, N., Holmes, I., Schlötterer, C. & Kosiol, C. Estimating empirical codon hidden Markov models. Molecular Biology and Evolution 30, 725–736 (2013).

31. De Maio, N. et al. Mutation rates and selection on synonymous mutations in SARS-CoV-2. Genome Biology and Evolution 13, evab087 (2021).

32. Sanderson, T. et al. A molnupiravir-associated mutational signature in global SARS-CoV-2 genomes. Nature 623, 594–600 (2023).

33. Markov, P. V. et al. The evolution of SARS-CoV-2. Nature Reviews Microbiology 21, 361–379 (2023).

34. De Maio, N. et al. Rate variation and recurrent sequence errors in pandemic-scale phylogenetics. bioRxiv. https://www.biorxiv.org/content/early/2024/07/15/2024.07.12.603240 (2024).

35. Hunt, M. et al. Addressing pandemic-wide systematic errors in the SARS-CoV-2 phylogeny. bioRxiv. 10.1101/2024.04.29.591666 (2024).

36. Finkel, Y. et al. The coding capacity of SARS-CoV-2. Nature 589, 125–130 (2021).

37. De Maio, N., et al. Issues with SARS-CoV-2 sequencing data. virolog-ical.org. https://virological.org/t/issues-with-sars-cov-2-sequencing-data/473 (2020).

38. Turakhia, Y., et al. Stability of SARS-CoV-2 phylogenies. PLoS Genetics 16, e1009175 (2020).

39. Smith, K., Ye, C. & Turakhia, Y. Tracking and curating putative SARS-CoV-2 recombinants with RIVET. Bioinformatics 39, btad538 (2023).

40. Lagerborg, K. A. et al. Synthetic DNA spike-ins (SDSIs) enable sample tracking and detection of inter-sample contamination in SARS-CoV-2 sequencing workflows. Nature Microbiology 7, 108–119 (2022).

41. Rockett, R. J. et al. Co-infection with SARS-CoV-2 Omicron and Delta variants revealed by genomic surveillance. Nature Communications 13, 2745 (2022).

42. Rambaut, A. et al. A dynamic nomenclature proposal for SARS-CoV-2 lineages to assist genomic epidemiology. Nature Microbiology 5, 1403– 1407 (2020).

43. O’Toole, Á. et al. Assignment of epidemiological lineages in an emerging pandemic using the pangolin tool. Virus Evolution 7, veab064 (2021).

44. Sanderson, T. Taxonium, a web-based tool for exploring large phylogenetic trees. Elife 11 (2022).

45. Turakhia, Y. et al. Pandemic-scale phylogenomics reveals the SARS-CoV-2 recombination landscape. Nature 609, 994–997 (2022).

46. V’kovski, P., Kratzel, A., Steiner, S., Stalder, H. & Thiel, V. Coronavirus biology and replication: implications for SARS-CoV-2. Nature Reviews Microbiology 19, 155–170 (2021).

47. Malone, B., Urakova, N., Snijder, E. J. & Campbell, E. A. Structures and functions of coronavirus replication–transcription complexes and their relevance for SARS-CoV-2 drug design. Nature Reviews Molecular Cell Biology 23, 21–39 (2022).

48. Bradley, C. C. et al. Targeted accurate RNA consensus sequencing (tARC-seq) reveals mechanisms of replication error affecting SARS-CoV-2 divergence. Nature Microbiology, 1–11 (2024).

49. Kim, D. et al. The architecture of SARS-CoV-2 transcriptome. Cell 181, 914–921 (2020).

50. Garushyants, S. K., Rogozin, I. B. & Koonin, E. V. Template switching and duplications in SARS-CoV-2 genomes give rise to insertion variants that merit monitoring. Communications Biology 4, 1343 (2021).

51. Chrisman, B. S. et al. Indels in SARS-CoV-2 occur at template-switching hotspots. BioData Mining 14, 1–16 (2021).

52. Liang, J. et al. How the replication and transcription complex functions in jumping transcription of SARS-CoV-2. Frontiers in Genetics 13, 904513 (2022).

53. Ugolini, C. et al. Nanopore ReCappable sequencing maps SARS-CoV-2 5*^′^* capping sites and provides new insights into the structure of sgRNAs. Nucleic Acids Research 50, 3475–3489 (2022).

54. Didelot, X. & Wilson, D. J. ClonalFrameML: efficient inference of recombination in whole bacterial genomes. PLoS Computational Biology 11, e1004041 (2015).

55. Yang, Z. & Nielsen, R. Codon-substitution models for detecting molecular adaptation at individual sites along specific lineages. Molecular Biology and Evolution 19, 908–917 (2002).

56. Nielsen, R. & Yang, Z. Likelihood models for detecting positively selected amino acid sites and applications to the HIV-1 envelope gene. Genetics 148, 929–936 (1998).

57. Zou, Z. & Zhang, J. Morphological and molecular convergences in mammalian phylogenetics. Nature Communications 7, 12758 (2016).

58. Pasternak, A. O., Spaan, W. J. & Snijder, E. J. Nidovirus transcription: how to make sense… ? Journal of General Virology 87, 1403–1421 (2006).

59. De Maio, N. Global (2M) SARS-CoV-2 genomes dataset, from Viridian, processed with MAPLE. Zenodo. 10.5281/zenodo.12733488 (2024).

60. Katoh, K., Misawa, K., Kuma, K.-i. & Miyata, T. MAFFT: a novel method for rapid multiple sequence alignment based on fast Fourier transform. Nucleic Acids Research 30, 3059–3066 (2002).

